# Role of Epigenetics in Unicellular to Multicellular Transition in *Dictyostelium*

**DOI:** 10.1101/2020.09.03.282152

**Authors:** Simon Yuan Wang, Elizabeth Ann Pollina, I-Hao Wang, Henry L. Bushnell, Ken Takashima, Colette Fritsche, George Sabin, Paul Lieberman Greer, Eric Lieberman Greer

## Abstract

The evolution of multicellularity is a critical event that remains incompletely understood. We use the social amoeba, *Dictyostelium discoideum,* one of the rare organisms that exists in both unicellular and multicellular stages, to examine the role of epigenetics in regulating multicellularity. While transitioning to multicellular states, patterns of H3K4 methylation and H3K27 acetylation significantly change. By combining transcriptomics, epigenomics, chromatin accessibility, and syntenic analyses with other unicellular and multicellular organisms, we identify 52 conserved genes, which are specifically accessible and expressed during multicellular states. We validated that four of these genes, including the H3K27 deacetylase *hdaD,* are necessary and that an SMC-like gene, *smcl1,* is sufficient for multicellularity. These results highlight the importance of epigenetics in reorganizing chromatin architecture to facilitate the evolution of multicellularity.

**One Sentence Summary:** Epigenetic regulation of multicellularity

## Main Text

A critical evolutionary event, the transition from unicellularity to multicellularity, requires the acquisition of cell-cell adhesion and cooperation, and frees the organism for growth and specialization (*1*). However, the molecular adaptations that facilitated this transition remain incompletely understood. Earlier studies proposed that the acquisition of new genes or new transcription factor networks drove the evolution of multicellularity (*2*). However, the emergence of multicellularity does not appear to be explained exclusively by the appearance of new genes as multicellular and unicellular genomes contain largely the same genes. Likewise the acquisition of new transcription factor networks does not always correlate with multicellularity (*3*). These observations suggest a potential role for chromatin regulation in the transition to multicellularity. Although many of the individual components for modern epigenetic regulatory systems are present in more ancient ancestors, their integration into a coherent system of gene regulation did not occur until eukaryotes. Moreover, epigenetic diversity expanded rapidly upon the transition to multicellularity (*4–7*). However, whether the increased complexity of epigenetic regulation may have helped to drive the transition to multicellularity is unknown.

We sought to address this question using *Dictyostelium discoideum* because it is one of the rare organisms that exists in both unicellular and multicellular states. When bacterial food is plentiful, *Dictyostelium* are single-celled amoebae. However, when the food supply is exhausted, *Dictyostelium* produce a chemoattractant, which signals to neighboring cells to aggregate into a multicellular organism of ~215,000 cells (*8, 9*). Within 22 hours, unicellular *Dictyostelium* aggregate into a multicellular mound that develops to a migrating slug state (*10*). After locating an appropriate environment, the slug will subsequently develop into a mature fruiting body, complete with both a stalk and spores that disseminate to distant locations (Fig. 1a). Once food becomes available, spores quickly germinate into single-celled amoebae (*11*). The rapidity with which this organism undergoes such dramatic physiological changes, while maintaining the same DNA code, make it an ideal model organism for studying the role of epigenetics in multicellularity. Additionally, *Dictyostelium* has a relatively small genome (34 megabases (*12*)), is easy to culture in a lab and to manipulate genetically (*13–15*) and can be induced to develop using simple environmental manipulations (*11*). Importantly, *Dictyostelium* contain the conserved epigenetic regulating systems of other eukaryotes (*16*), including histone modifications and modifying enzymes that regulate gene expression (*17–19*).

**Fig. 1.**
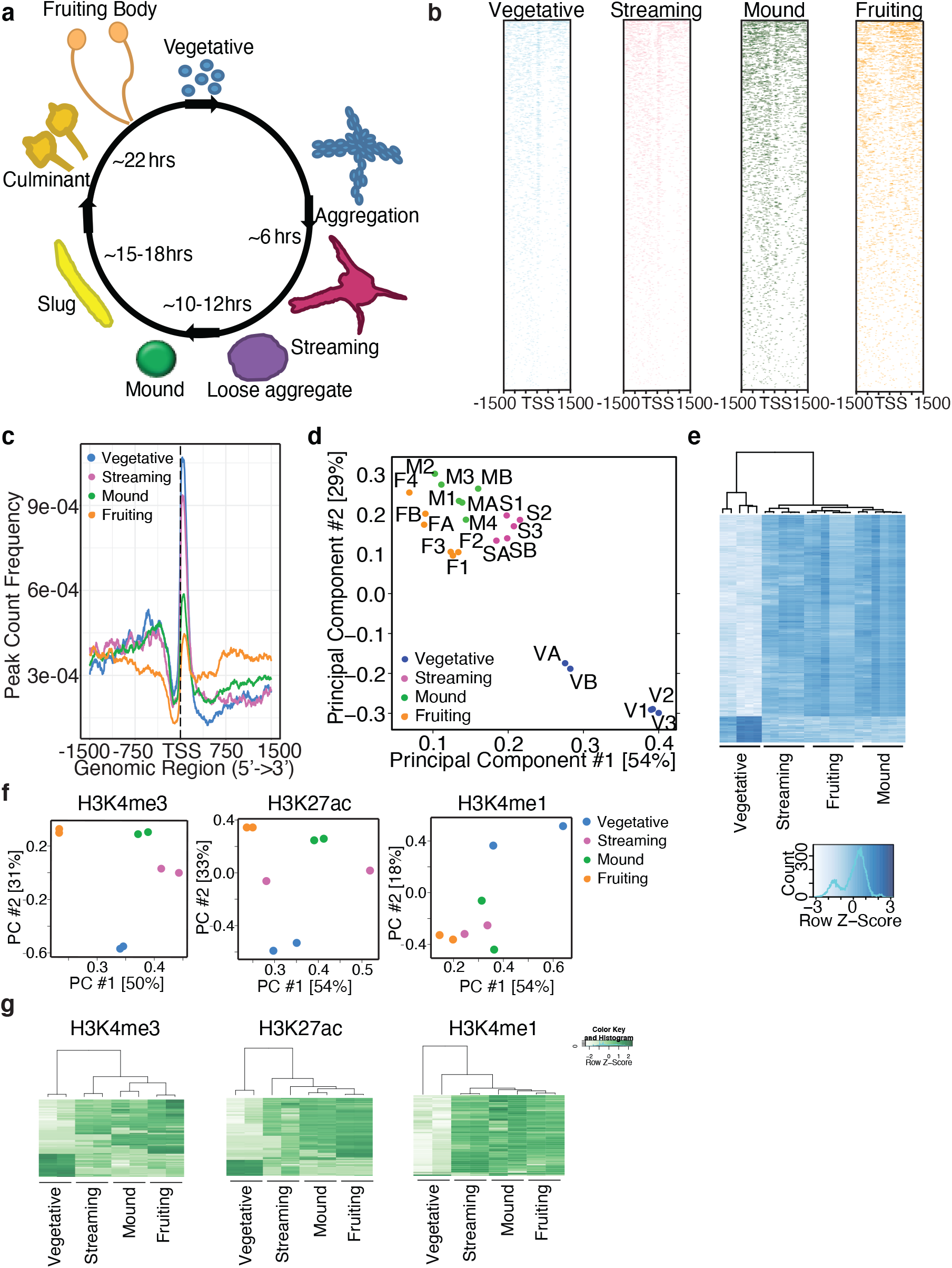
Chromatin modifications and accessibility on *D. discoideum* display stage-specific patterns for H3K4me3, H3K27ac, H3K4me1, and chromatin architecture. **a**, Cartoon diagram representing the different developmental stages and approximate time required to achieve each stage. Colors used in this diagram are used throughout the figures to represent specific stages. **b**, Heatmaps of ATAC-seq peaks on each of the stages of *Dictyostelium* 1500 basepairs upstream or downstream of the TSS. **c**, Metagene analysis of ATAC-seq experiments performed at different developmental stages shows that the unicellular stage of *D. discoideum* (vegetative: shown in blue line) displays greater chromatin accessibility 250 basepairs downstream of the TSS than multicellular stages (streaming: shown in purple line, mound: shown in green line, fruiting body: shown in orange line). **d**, PCA analysis shows the open regions in vegetative *D. discoideum* are separate from those in the streaming, mound, and fruiting stages. A and B mark replicates from frozen samples while 1, 2, 3, and 4 mark replicates from fresh samples. **e**, A heatmap of the ATACseq data at different life cycle stages shows distinct accessible regions in the multicellular stages relative to the unicellular stage. **f,** PCA of ChIPseq analyses of H3K4me3, H3K27ac, and H3K4me1 reveals that H3K4me3, H3K27ac, and H3K4me1 are sufficient to distinguish unicellular from multicellular *D. discoideum*. **g**, Heatmaps show distinct chromatin modification patterns for the localization of H3K4me3, H3K27ac, and H3K4me1 in unicellular (vegetative) and multicellular (streaming, mound, and fruiting body) stages. H3K36me3 did not display distinct patterning (Extended Data Fig. 6d) and H3K27me3, H3K9me3, and H3 were unable to produce heat maps due to either poor antibody specificity or similar localization occurrences.

Here we provide the first comprehensive map of epigenetic states (RNA expression, histone modifications, and chromatin accessibility) across four life cycle stages of *Dictyostelium*. By comparing 2473 syntenic gene expression patterns and chromatin accessibility between unicellular and multicellular *Dictyostelium* with that of fission yeast *S. pombe* and *C. elegans*, we identify candidate genes that could play important roles in multicellular development. Knockout of four multicellular genes caused delays in multicellularity while overexpression of one of these multicellular genes increased multicellularity. Our findings identify critical, novel regulators of multicellularity, and reveal that epigenomic changes correlate with the transition from unicellularity to multicellularity.

## Results

### *Dictyostelium* display unicellular and multicellular specific chromatin patterns

To begin to determine whether chromatin modifications might contribute to *D. discoideum* life cycle changes, overall histone methylation and acetylation levels were assessed by western blots at different developmental stages. These analyses revealed that H3K4me3 levels decreased and H3K4me1 increased significantly upon the transition to multicellularity suggesting that chromatin modification might help regulate the transition to multicellularity in *Dictyostelium* (Extended Data Fig. 1). To more precisely ascertain where within the genome specific changes in chromatin state occur during this transition, we performed Assay for Transposase-Accessible Chromatin using sequencing (ATACseq) (*20*), which broadly identifies which regions of the chromatin are open and accessible for transcription and which regions are closed and quiescent at four stages of *D. discoideum* life cycle: the vegetative, streaming, mound, and fruiting stages. We performed ATACseq on both frozen and fresh samples with highly reproducible results (Extended Data Fig. 2). Similar to most eukaryotes (*21*), the chromatin is compacted directly preceding the transcriptional start sites (TSS) (Fig. 1b). By performing a metagene analysis of the ATACseq data, we found that there was a decrease in accessibility immediately after the TSS as *Dictyotelium* transition from the unicellular to multicellular forms (Fig. 1c and Extended Data Fig. 3a). A PCA and clustering analyses of the ATACseq datasets clearly separated the vegetative unicellular stage from the multicellular stages, indicating that the accessible chromatin might reveal genomic patterns unique to unicellular and multicellular states of *D. discoideum* (Fig. 1, d and e, and Table S1). This was accompanied by a progressive increase of accessibility at promoters and a progressive increase of compaction at exons as *Dictyostelium* proceed from vegetative toward fruiting body stage (Extended Data Fig. 3b). A motif analysis of accessible regions revealed a motif, CCATATATGG, that was enriched in all multicellular stages but was absent in the unicellular stage (Extended Data Fig. 4). This motif provides a binding site for MADS-box transcription factors (*22*), which, interestingly, have been implicated in the evolution of multicellularity (*23*). Deletion of two *Dictyostelium* MADS-box transcription factors, *mef2A* and *srfA,* has been shown to slow developmental progression (*24–26*). These results raise the possibility that to transition to multicellularity, *Dictyostelium* reorganizes its chromatin structure to make binding sites for multicellular specific transcription factors *mef2A* and *srfA* accessible for binding, likely to facilitate the transcription of their target genes. Together these data reveal unicellular and multicellular *D. discoideum* have different patterns of chromatin accessibility regions throughout the genome, which could expose different transcription factor binding sites at specific stages.

To characterize chromatin modifications, which might explain these alterations in chromatin accessibility, ChIP-seq was performed in duplicate at the four stages analyzed by ATACseq using antibodies that recognize Histone H3, Histone H3 lysine 4 monomethylation (H3K4me1), H3K4 trimethylation (H3K4me3), H3K9me3, H3K27me3, H3K36me3, or H3K27 acetylation (H3K27ac) (Fig. 1, f-g and Extended Data Figs. 5 and 6). A high correlation in ChIPseq peaks was observed across biological replicates indicating good specificity and reproducibility (Extended Data Fig. 5). A principal component analysis (PCA) of observed ChIPseq patterns revealed that certain chromatin modifications accurately distinguished between the life cycle stages of *Dictyostelium*. For instance, H3K4me3 and H3K27ac marks clearly separated the unicellular and multicellular stages from each other (Fig. 1, f and g). H3K4me1 modifications were able to separate the unicellular stage from the multicellular stages but were not as powerful at distinguishing the different multicellular stages (Fig. 1, f and g). H3K36me3 had some capacity to distinguish stages at the 2^nd^ principal component, but H3K27me3, H3K9me3, and Histone H3 did not discern distinct stages even by the 3^rd^ principal component (Extended Data Fig. 6b, and Tables S2 and S3), suggesting that these marks may not change consistently with life cycle stage. The chromatin modifications that were capable of distinguishing unicellular from multicellular stages were similar to those marks that we observed to have different overall levels across the life cycle of *Dictyostelium* (Extended Data Fig. 1). Additionally, chromatin marks with the capacity to distinguish stages correlate with gene activation, while marks that correlate with gene repression are unable to distinguish *Dictyostelium* stages. This observation suggests that activating chromatin marks may be more important for determining cellular states than repressive marks. Together, these data demonstrate that chromatin modifications change as *D. discoideum* transitions through different life cycle stages and that the most dramatic chromatin modification changes occur in H3K4 methylation and H3K27 acetylation upon the transition from unicellular to multicellular states raising the possibility that regulation of these specific chromatin marks could play a critical role in the transition to multicellularity.

### Histone deacetylation and methylation inhibitors delay development

To begin to test whether changes in H3K4 methylation and H3K27 acetylation affect the transition to multicellularity, we treated vegetative *D. discoideum* with chemical inhibitors of histone deacetylation and methylation enzymes and then induced multicellularity by starvation. The histone deacetylation inhibitor Trichostatin A (TSA), the histone methylation enzyme inhibitors sinefungin and BIX 01294 all caused a concentration dependent delay in the initiation of multicellularity, defined by the amount of time required to achieve the mound stage, as well as a delay in the completion of multicellular development, defined by the time required to produce fruiting bodies (Extended Data Fig. 7). We confirmed that TSA caused an increase in H3K27ac while sinefungin caused a decrease in H3K4me3 and H3K9me3. BIX01294, although reported to be an H3K9me3 inhibitor, led to a decrease in H3K4me3 and H3K27ac without any measurable effects on H3K9me3 in 293T cells (Extended Data Fig. 7a). These inhibitors delayed the initiation of multicellularity from ~12 hours in control conditions to ~17 hours in 5 μM sinefungin (p=0.001), ~15 hours in 10 μM BIX 01294 (p=0.047), and ~17 hours in 1 μM TSA (p=0.0024) (Extended Data Fig. 7, b and c). Histone modification chemical inhibition similarly delayed the completion of multicellular development (Extended Data Fig. S7, b and c). While *D. discoideum* took ~22 hours to reach the fruiting body stage under basal conditions, 10 μM BIX 01294, 5 μM sinefungin, and 1 μM TSA significantly increased this time to 27, 37, and 29 hours, respectively (Extended Data Fig. 7, b and c, BIX 01294 p=0.0126; sinefungin p<0.0001; TSA p=0.0009). Treatment with sinefungin also caused a reduction in the number of cells making up the multicellular *D. discoideum* (Extended Data Fig. 7d). The diameter of the fruiting body, which correlates with the number of cells composing the multicellular form (*8, 9*), was ~9 μm without treatment and ~6 um after 5 μM sinefungin treatment (p<0.05). To determine whether these inhibitors specifically delayed multicellularity or were more generally toxic, the effects of the lowest concentrations of TSA, sinefungin, and BIX 01294 that delayed multicellularity were measured for effects on chemotaxis and apoptosis. Although 1 μM TSA and 5 μM sinefungin significantly reduced chemotaxis towards folate (Extended Data Fig. 8, a and b, p<0.0001 and p=007, respectively), 10 μM BIX 01294 had no significant effect on chemotaxis (Extended Data Fig. 8, a and b, p=0.9978). Additionally, 10 μM BIX 01294 and 5 μM sinefungin had no effect on apoptosis as assessed by trypan blue staining (Extended Data Fig. 8c), propidium iodide (PI) staining (Extended Data Fig. 8d), or annexin V staining (Extended Data Fig. 8e) while 1 μM TSA caused a modest increase in apoptosis when assessed by annexin V staining (Extended Data Fig. 8e). Thus, chemicals which regulate H3K4 methylation inhibit multicellularity without having dramatic effects on other properties of *D. discoideum*. Confirming the importance of H3K4 methylation in multicellularity, previous work has demonstrated that deletion of the H3K4me3 methyltransferase *set1*, causes unusually rapid development (*18*). Taken together, these results suggest that changes to the amounts or locations of chromatin modifications might regulate multicellularity.

### Stage-specific gene expression patterns

To begin to investigate whether the chromatin changes we observed correlate with gene expression changes, we performed RNA-seq analysis on vegetative, streaming, mound, and fruiting stages of *D. discoideum*. Biological replicates were reproducible (Extended Data Fig. 9a), individual stages were readily distinguished from each other by correlation analysis in a heatmap and Bland-Altman plots (Extended Data Fig. 9, b-f), and our results correlated well with previously generated RNAseq data (Extended Data Fig. 9, g-j) (*27*). As expected, the expression of some genes remained constant, some declined, and others increased at each transition (Fig. 2a and Table S4). A principal component analysis (PCA) readily distinguished individual stages. The developmentally similar mound and streaming stages clustered close to each other, while the unicellular vegetative and the final fruiting body stage clustered far from each other (Fig. 2b). Importantly, this approach recapitulates previously reported stage specific patterns of gene expression (*27*), (Fig. 2c). We confirmed by qRT-PCR the stage-specific gene expression patterns of 5 multicellular and 1 unicellular enriched genes (Fig. 2d). Together, these data suggest that the RNAseq analysis is reliable. Gene ontology (GO) analysis revealed stage-specific enrichment of gene expression in distinct processes and pathways (Fig. 2e). Using a threshold FDR of < 0.05, 1216 genes were enriched in the unicellular state and 1649 genes were enriched in the multicellular states (Fig. 2f). By GO analysis, unicellular genes were enriched for genes involved in protein synthesis (Fig. 2g, upper panel and Table S4). Conversely, multicellular enriched genes were selectively involved in cell differentiation, development, and adhesion and cellular components required for extracellular elements, such as the cell surface and the plasma membrane (Fig. 2g, lower panel and Table S4). These changes reflect the shifting cellular priorities during multicellular stages.

**Fig. 2.**
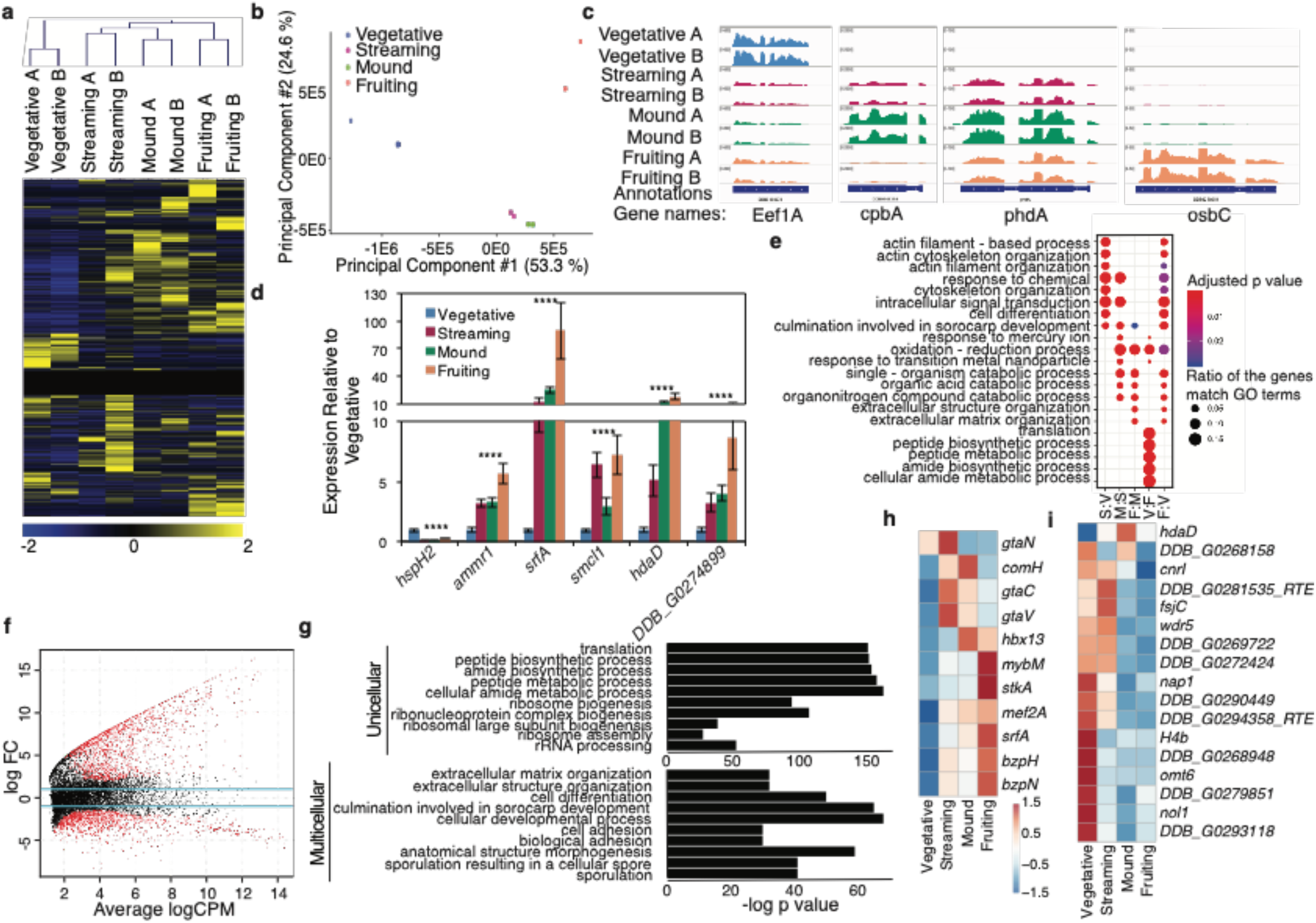
Transcriptomic analysis on vegetative, streaming, mound, and fruiting body stages of *D. disoideum* reveals unicellular and multicellular specific gene expression patterns. **a**, A heatmap of gene expression of all 13,267 genes at different stages of *D. discoideum* reveals gene sets that increase, decrease, and remain constant throughout development. Scale bar is displayed below. A and B indicate independent replicates. **b**, Principle components analysis (PCA) of gene expression demonstrates that biological replicates cluster together and unicellular, vegetative stage clusters separately from the three multicellular stages. **c**, Representative stage-specific gene expression tracks of known stage-specific genes. **d**, Real-time qPCR validation of representative multicellular and unicellular gene expression from RNAseq datasets. This graph represents the mean ± the standard deviation of three biological replicates performed in triplicate. **e**, Stage by stage Gene Ontology (GO) comparison of differentially regulated genes reveals that classes of genes which correlate with multicellularity are misregulated when comparing Vegetative (V) to Streaming (S) or Fruiting body (F) stages. **f**, A Bland-Altman plot of *D. discoideum* RNAseq datasets reveals genes which are upregulated and downregulated between unicellular and both multicellular stages. Red dots represent significantly differentially expressed genes, black dots represent not significantly differentially expressed genes. **g**, The top ten GO terms of unicellular enriched (top) or multicellular enriched (bottom) processes reveals genes involved in essential processes are enriched in unicellular *D. discoideum* and processes required for development, differentiation, communication, and adhesion are enriched in multicellular *D. discoideum.* Heat maps of transcription factors **h**, and chromatin modifying enzymes **i**, that display unicellular or multicellular enriched expression are displayed here. An analysis of all transcription factors and all chromatin modifying enzymes identified in *D. discoideum* is presented in Extended Data Fig. 10.

As transcription factors play an important role in the transition to multicellularity (*28, 29*), we next identified transcription factors that had stage-specific expression (Extended Data Fig. 10a). Ten transcription factors were expressed predominantly in multicellular stages, but none was found to be predominant in the vegetative stage (Fig. 2h). Two of these transcription factors, were the MADS-box transcription factors *srfA* and *mef2A,* which were previously shown to play a role in *D. discoideum* development (*24–26*). We found that genes which were upregulated in multicellular states were enriched for MADS-box transcription factor binding motifs (Extended Data Fig. 10c p = 4.08e-25). These results suggest that not only does the chromatin reorganize to expose *srfA* and *mef2A* binding sites (Extended Data Fig. 4), but *srfA* and *mef2A* expression is coordinately upregulated (Fig. 2, d and h) to facilitate the transcription of SrfA and Mef2A targets.

We also examined stage specific expression of all putative chromatin modifying and binding proteins, identified based on homology (Extended Data Fig. 10b). 17 chromatin modifying enzymes or binding proteins displayed unicellular or multicellular specific expression patterns (Fig. 2i). More transcription factors whose expression varied with developmental stage were upregulated in the multicellular stages, whereas more chromatin modifying enzymes were upregulated in the unicellular stage (Extended Data Fig. 10, a and b, *p* < 0.05). Notably, one of the chromatin regulating enzymes, the putative histone deacetylase *hdaD,* was selectively enriched in multicellular states (Fig. 2, d and i), raising the possibility that *hdaD* could be activated in multicellular states to deacetylate histones and reorganize chromatin to facilitate the transition to multicellularity.

### Comparative epigenomic analysis of *S. pombe,* unicellular and multicellular *D. discoideum*, and *C. elegans* identifies unicellular and multicellular signatures

To begin to determine whether chromatin modifications or accessibility could be used to identify critical regions whose chromatin changed upon multicellularity, we first investigated whether chromatin modifications in *D. discoideum* are interpreted in a similar manner as they are in other eukaryotes. We therefore characterized how gene expression correlated with the ChIP-seq and ATACseq data we generated in *D. discoideum*. As has previously been demonstrated in mammalian cell lines (*30, 31*), chromatin accessibility correlated with gene expression in *Dictyostelium*; the top 1000 most lowly and highly expressed genes displayed lower and higher chromatin accessibility, respectively (Fig. 3a and Table S5). Histone modifications of specific genes also correlated with their expression in similar ways in *D. discoideum* as they do in other eukaryotic organisms. For instance, *srfA,* which turns on during multicellular stages (Fig. 2d), exhibited increased H3K4me3, decreased H3K27me3, and increased chromatin accessibility at its promoter at multicellular stages relative to the unicellular stage, which is when it is more highly expressed (Fig. 3b).

**Fig. 3.**
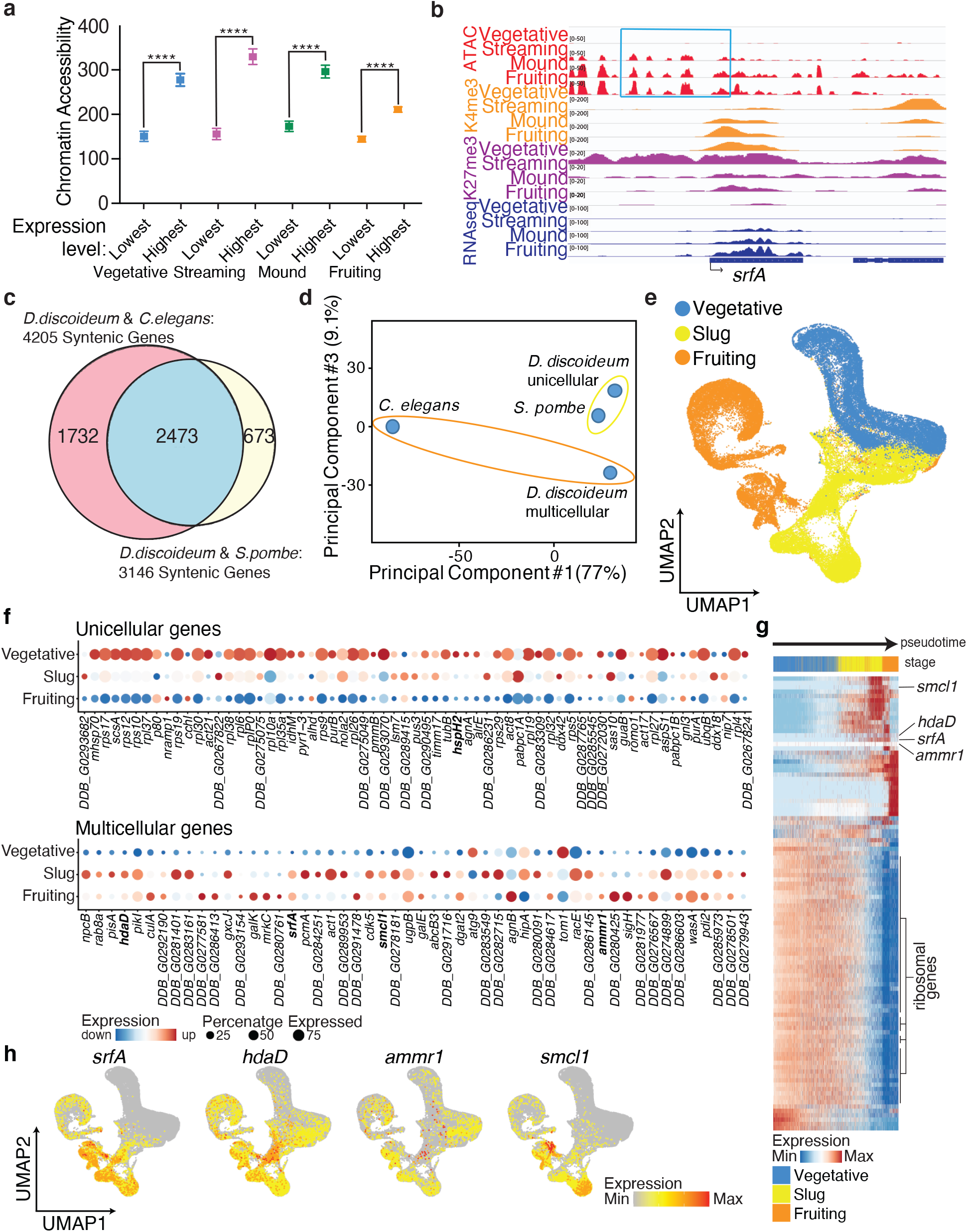
Comparative epigenomic analysis of *S. pombe,* unicellular and multicellular *D. discoideum*, and *C. elegans* identifies unicellular and multicellular signatures. **a**, The 1000 most highly expressed genes at each life cycle stage display a higher degree of chromatin accessibility than the 1000 most lowly expressed genes at each life cycle stage. ****:p<0.0001 as assessed by one-way ANOVA. **b**, Representative integrative genomics viewer (IGV) tracks of H3K4me3 (orange), H3K27me3 (purple), ATAC-seq (red), and gene expression (navy) at *srfA* demonstrates increases in H3K4me3, decreases in H3K27me3, and increased chromatin accessibility at the promoters correlate with increases in gene expression of *D. discoideum* at four life cycle stages. **c**, A Venn diagram displays the 2473 conserved genes in *S. pombe, D. discoideum,* and *C. elegans.* **d**, PCA of syntenic genes illustrates that unicellular enriched gene expressions of *D. discoideum* cluster closer to *S. pombe* than to multicellular *D. discoideum* by the third principal component. **e**, Single cell RNA sequencing analysis of vegetative (blue), slug (yellow), and fruiting body (orange) cells reveals unique gene expression signatures of different stage *D. discoideum.* Uniform manifold approximation and projection (UMAP) is displayed for visual representation. **f**, Single cell RNAseq expression correlates highly with bulk RNAseq expression as exemplified by 67 “unicellular” genes and 52 “multicellular” genes. The 67 genes represented are accessible and expressed in unicellular *D. discoideum* and *S. pombe* but not in multicellular *D. discoideum* and *C. elegans* and the 52 genes represented display the opposite accessibility and expression as identified from syntenic analysis of bulk RNAseq and ATACseq datasets. Highlighted in bold are the genes which we knocked out in this study. **g**, The 100 genes whose expression changes allow for the quantitative measurement of progression of *D. discoideum* from vegetative through slug to fruiting body stages are displayed in this heatmap pseudotime analysis. Expression of *srfA, hdaD, ammr1,* and *smcl1* are also included in this heatmap. Ribosomal gene expression plays an outsized role in determining pseudotime and these genes are marked. **h**, Representative UMAP graphs display the expression levels of *srfA, hdaD, ammr1,* and *smcl1* from single cell RNAseq analysis.

To investigate further whether epigenomic changes could correlate with unicellular or multicellular states, we compared the epigenomic signatures of unicellular and multicellular *D. discoideum* to two other eukaryotes whose epigenomes are well defined, the unicellular *Schizosaccharomyces pombe* and the multicellular *Caenorhabditis elegans,* which are both ~1480 million years evolutionarily separated from *D. discoideum* (*32, 33*). We identified 2473 conserved regions by predicting orthology and paralogy relationships in all three species by metaphors using pre-computed phylogenetic trees (*34*) (Fig. 3c and Table S6). The majority of these highly conserved genes (79.1%) were evenly expressed across all *Dictyostelium* stages while 15.6% were unicellular enriched and 5% were multicellular enriched (Extended Data Fig. 11). Interestingly, a PCA analysis of the syntenic genes showed that by principal component 3 the expression patterns of unicellular *Dictyostelium* genes were more similar to that of *S. pombe* than to multicellular *Dictyostelium* genes (Fig. 3d).

To determine whether epigenetic features could underlie these unicellular and multicellular gene expression signatures, we examined whether accessible chromatin regions correlated with unicellular or multicellular signatures. We examined the ATACseq profiles of *C. elegans* (*35*) and *S. pombe* (*36*) at the 2473 evolutionarily conserved regions of the genome and found that of the conserved genes, 1352 and 804 were accessible, respectively. Of the *D. discoideum* conserved genes, 922 were accessible in the unicellular stage while 720 were accessible in the multicellular stages. Among those 922 accessible regions in unicellular *D. discoideum*, 132 were accessible in multicellular *Dictyostelium* or *C. elegans* (14.3%), but 289 sites were exclusively accessible in *S. pombe* (31.3%). Similarly, among the 720 accessible regions in multicellular *D. discoideum*, 94 were accessible in unicellular *Dictyostelium* or *S. pombe* (13.1%), but 333 sites were exclusively accessible in *C. elegans* (46.3%). We further reduced these lists of 289 unicellular accessible and 333 multicellular accessible regions to genes which were expressed exclusively in unicellular or multicellular stages. This analysis left us with 67 genes which were accessible and expressed in unicellular state or organisms and inaccessible and not expressed in multicellular state or organisms and 52 genes which were accessible and expressed in multicellular state or organisms and inaccessible and not expressed in unicellular state or organisms (Table S7), suggesting that these accessible chromatin regions correlate with unicellularity or multicellularity, respectively across evolution. Interestingly, *srfA,* the MADS-box transcription factor that had previously been shown to play a role in development in *D. discoideum* (*24*), was one of the genes present in the multicellular list suggesting that our method of identifying critical genes involved in multicellularity based on their chromatin states was successful.

Because bulk RNAseq experiments do not guarantee that all cells are at the precise stage, and changes in gene expression could be masked by cells that are not critical for transition states, we performed single-cell RNAseq experiments. This analysis clearly distinguished the unicellular state from the multicellular stages (Fig. 3e). We found that the expression of the 67 unicellular genes and 52 multicellular genes, identified based on the syntenic analysis of their expression in bulk RNAseq and ATACseq experiments, correlated well with the single cell RNAseq experiments (Fig. 3f). We next used the single cell RNAseq data to examine which genes’ expression signatures could account for the progression of *D. discoideum* from vegetative through slug to fruiting body stages. This lineage tracing analysis revealed that of the top 100 genes whose expression changes helped determine this temporal progression, 54 of them are ribosomal genes (Fig. 3g). This finding demonstrates that the ribosomal heterogeneity that was initially discovered and reported in *Dictyostelium* (*37*) is predominantly driven by gene expression changes in ribosomal protein genes. Interestingly, 13 of these ribosomal protein genes were identified as part of our unicellular signature (Fig. 3f and Table S7), indicating that changes in chromatin accessibility at ribosomal protein genes could be a critical driver of the evolution of multicellularity. We further confirmed that 4 genes that were accessible and expressed in *C. elegans* and multicellular *Dictyostelium* and quiescent in unicellular state or *S. pombe* were highly enriched or only expressed in multicellular cells (Fig. 3h).

### Deletion of epigenomically identified multicellular genes delays or prevents multicellularity in *D. discoideum*

Together, these approaches identified a total of 67 “unicellular” genes, which were accessible and expressed in unicellular state or organisms and inaccessible and not expressed in multicellular state or organisms and 52 “multicellular” genes, which were accessible and expressed in multicellular state or organisms and inaccessible and not expressed in unicellular state or organisms. To begin to test whether these identified genes contribute to the transition between unicellularity and multicelluarity, we analyzed knockouts of a subset of these genes. We obtained a previously generated knock-out strain of *srfA* (*38*) and attempted to delete 4 additional “multicellular” genes and 3 “unicellular” genes and examined their effects on rates of multicellularity. Despite repeated attempts, one “multicellular” gene (*DDB_G0274899*) and two “unicellular” gene (*DDB_G0267824* and *DDB_G0277333*) knock-outs were inviable, suggesting that these genes are essential for *D. discoideum* survival. Successful knock-outs of the remaining 4 genes were validated by sequencing (Extended Data Fig. 12).

Deletion of all four “multicellular” genes: *srfA; hdaD*; *DDB_G0283437*, a gene which we renamed *ammr1* based on its homology to the uncharacterized Alport syndrome gene Ammerc1; and *DDB_G0288013*; a gene which we renamed *smcl1* based on its similarity to a structural maintenance of chromatin (SMC) genes, delayed the transition to multicellularity and extended the entire developmental progression (Fig. 4, a and b: *srfA* p=0.0014 and p<0.0001; *hdaD* p=0.0165 and p=0.0011; *ammr1* p<0.0001 and p<0.0001; *smcl1* p <0.0001 and p<0.0001 respectively). For instance, knock out of *hdaD* caused the initiation of multicellularity to be delayed from 10.75 hours in control cells to 13.6 hours and the time required to achieve a fruiting body to be delayed from 22.11 hours to 29.88 hours (Fig. 4, a and b). Similarly, *srfA* knock-out caused a delay of the initiation of multicellularity (16.96 hours, p=0.0022) and a delay in complete development (38.83 hours, p<0.0001) and a reduction in the number of cells in the multicellular state to 45% of the wildtype *Dictyostelium* (Fig. 4c, p=0.0405). Likewise, deletion of *ammr1* dramatically slowed multicellularity to 45.8 hours (p<0.0001) and caused complete development to take 87.33 hours (p<0.0001), while deletion of *smcl1* caused a complete blockage in the progression past the aggregation stage to any multicellular stages (Fig. 4, a and b). By contrast, knock-out of the “unicellular” gene *DDB_G0293674,* a gene which we renamed *hspH2* based on its homology to the heat shock protein hsc70-3, had no effect on the transition to multicellularity or the developmental progression (Fig. 4, a and b, p=0.5166 and p=0.2585 respectively).

**Fig. 4.**
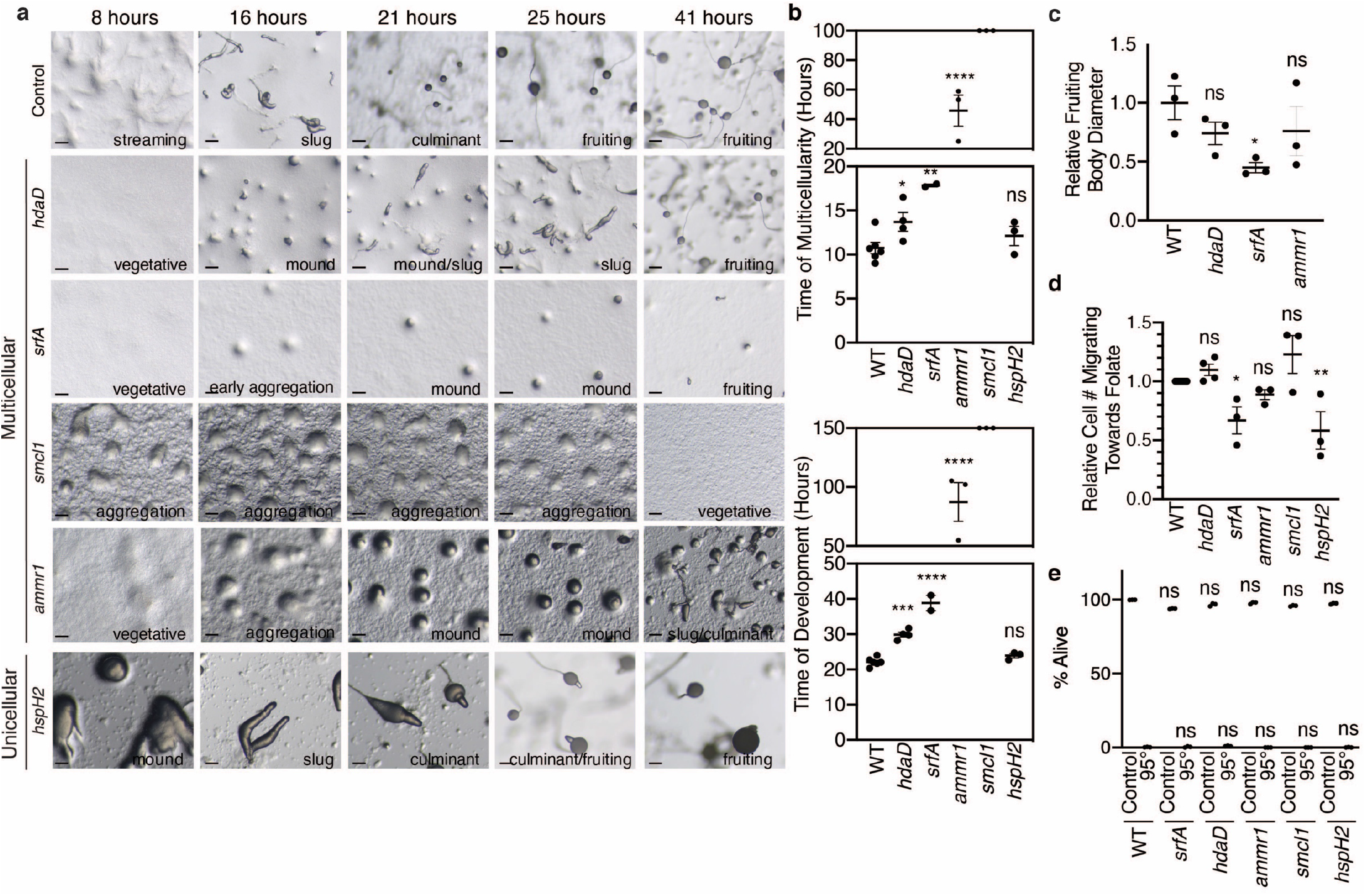
Genes identified as accessible and expressed in multicellular but not unicellular stages and organisms are necessary for multicellularity in *D. discoideum*. **a**, Representative images illustrating that knock-out of the multicellular genes, *hdaD, srfA, ammr1,* and *smcl1,* but not the unicellular gene, *hspH2,* results in developmental delays. Representative images are shown at 8, 16, 21, 25, and 41 hours and the developmental stage is noted in the bottom right hand corner of each image. Scale bars are 20 microns. **b**, Knockout of multicellular genes delays multicellularity (upper graph) and total development (lower graph) but knockout of a unicellular gene does not. This graph represents the mean ± the standard error of the mean of three independent biological replicates performed in triplicate. Ns: not significant, *: p<0.05, **: p<0.005, ***: p<0.001, ****:p<0.0001 as assessed by multiple comparison one-way ANOVA analysis. **c**, Knockout of *srfA* but not *hdaD* or *ammr1* caused a reduction in the diameter of the fruiting body as a proxy for the number of cells in each multicellular organism. This graph represents the mean ± the standard error of the mean of three independent biological replicates performed triplicates. Ns: not significant, *; p<0.05 as assessed by multiple comparison one-way ANOVA analysis. **d**, Knock-out of multicellular genes: *hdaD, ammr1,* and *smcl1* had no effect on chemotaxis while knock-out of *srfA* and unicellular gene *hspH2* decrease the chemotaxis capacity of *D. discoideum.* The bar graph represents the mean ± the standard error of the mean of three biological replicates performed in triplicate. The number of cells which migrated towards the 30° segment containing the location where 250 μM folate was placed were counted. Ns: not significant, *: p<0.05, **: p<0.005 as assessed by multiple comparison one-way ANOVA analysis. **e**, Knock-out of unicellular and multicellular genes had no effect on apoptosis as assessed by PI staining. All knock-out strains responded similarly to the WT strain in response to 95°C heat shock for 50 seconds. Each bar represents the mean ± the standard deviation of three biological replicates. Ns: not significant as assessed by one-way ANOVA analysis.

To determine whether deletion of these “multicellular” genes was simply reducing overall fitness, or was more specifically affecting *D. discoideum* multicellularity and development, we examined the effect of gene knock-outs on chemotaxis and apoptosis. While the “unicellular” gene *hspH2* displayed no effect on *D. discoideum* development, it did result in chemotaxis delay (Fig. 4d, p<0.0001), suggesting that this gene is essential for unicellular phenotypes. Of the “multicellular” genes only *srfA* displayed a decrease in chemotaxis (p=0.0144 by one-way ANOVA) while *hdaD, ammr1,* and *smcl1* were indistinguishable from control cells (Fig. 4d, p=0.7128, p=.06836, and p=0.1199, respectively). Additionally, knock-outs of all “multicellular” genes had no effect on apoptosis under basal conditions or in response to a 95°C heat shock as assessed by PI staining (Fig. 4d), suggesting that these deletions are not simply making the *Dictyostelium* sick and are more specifically affecting the transition to multicellularity.

To begin to determine whether the function of the genes that we identified as being necessary for multicellularity were also sufficient to induce a multicellular state, we overexpressed two of them, *hdaD* and *smcl1,* in WT *Dictyostelium.* We confirmed the overexpression of these genes by real-time RT PCR (Extended Data Fig. 13a). While overexpression of neither *hdaD* nor *smcl1* accelerated the timing of multicellularity (in fact *smcl1* overexpression caused a slowing of multicellularity timing) (Fig. 5, a and b and Extended Data Fig. 13, b and c), overexpression of *smcl1* did cause a ~3 fold increase in the number of cells which make up a multicellular *Dictyostelium* organism (Fig. 5, c-e), suggesting that *smcl1* enhances multicellularity. We found that overexpression of *smcl1* had no effect on chemotaxis (Fig. 5f, p=0.1207). Together these findings suggest that a gene which we identified based on its epigenomic status, the SMC like gene *smcl1,* was both necessary and sufficient to regulate multicellularity.

**Fig. 5.**
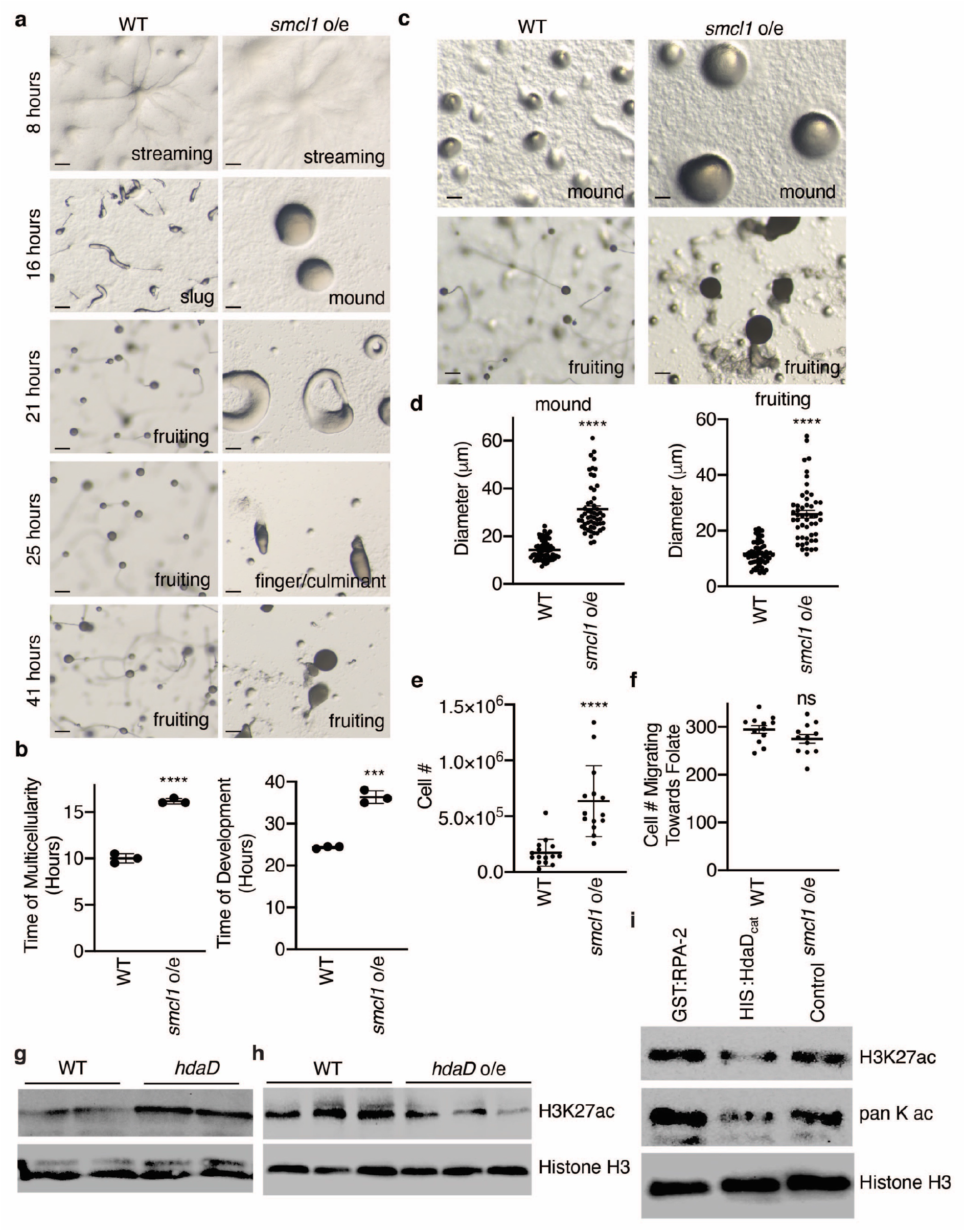
*smcl1* overexpression is sufficient for multicellularity in *D. discoideum* and HdaD is an H3K27 deacetylase. **a**, Representative pictures of *D. discoideum* development on KK2 plates show developmental delay after overexpression of *smcl1* at 8, 16, 21, 25 and 41 hours. Stage is displayed in the lower right hand corner. Scale bars are 20 microns. **b**, Overexpression of *smcl1* delays *D. discoideum* development, time to mound stage is shown in graph on top and complete development shown in graph on bottom. Each column represents the mean ± the standard error of the mean of three biological replicates performed in triplicate. ***: p<0.0005, ****: p<0.0001, as assessed by unpaired t test. **c**, Representative pictures of *D. discoideum* show increased size of mound (top) and fruiting body (bottom) after overexpression of *smcl1.* Scale bars are 20 microns. **d**, Overexpression of *smcl1* was sufficient to increase the size of mound (top) and fruiting body (bottom) diameter. Each column represents the mean ± the standard error of the mean of 54-67 *D. discoideum* multicellular organisms. ****: p<0.0001 as assessed by unpaired t test. **e**, Overexpression of *smcl1* was sufficient to increase the number of cells in each fruiting body. Each column represents the mean ± the standard error of the mean of 14-15 *D. discoideum* fruiting bodies. ****: p<0.0001 as assessed by unpaired t test. **f**, Overexpression of *smcl1* had no effect on chemotaxis of *D. discoideum.* Each column represents the mean ± the standard error of the mean of three biological replicates performed in triplicate. The number of cells which migrated towards the 30° segment containing 250 μM folate was counted. Ns: not significant, as assessed by unpaired t test. **g**, Representative western blots demonstrate an increase in H3K27ac in *hdaD* knock-out strains. Two independent biological replicates are displayed with the upper blot probed with H3K27ac and the bottom blot probed with Histone H3 antibodies. **h**, *hdaD* overexpression causes a decrease in H3K27ac as assessed by western blot. **i**, His-tagged HdaD catalytic domain (His:HdaD_catalytic_) but not GST-tagged RPA-2 is able to deacetylate lysine 27 of histones purified from *Dictyostelium.* This western blot is representative of 4 independent experiments.

Together, these experiments identified four genes that regulate the transition from unicellularity to multicellularity. To begin to investigate the mechanisms by which these genes control multicellularity, we focused on *hdaD*, which sequence alignment suggested might encode a histone deacetylase. HDACs are attractive candidates to mediate these processes because of their capacity to deacetylate histones and therefore cause a reorganization of the chromatin. Western blotting of *hdaD* knockouts revealed an increase in H3K27ac when compared to wild type *Dictyostelium* (Fig. 5g), suggesting that HdaD is a H3K27 deacetylase in *D. discoideum*. Consistent with this hypothesis, overexpression of *hdaD* caused a decrease in H3K27 acetylation (Fig. 5h). To determine whether HdaD is a direct H3K27 deacetylase we expressed a polyhistidine (His)-tagged catalytic domain of HdaD in bacteria, purified His-HdaD_catalytic domain_ to a single band, and analyzed its ability to deacetylate histones purified from *D. discoideum.* We found that His-HdaD_catalytic domain_, but not control proteins, was able to deacetylate H3K27 in purified histones (Fig. 5i), suggesting that HdaD is an H3K27 deacetylase *in vitro* and *in vivo*.

The finding that *hdaD* deletion inhibits multicellularity highlights the importance of H3K27 acetylation regulation that we had observed correlating on a global level with multicellularity in our ChIPseq experiments (Fig. 1, f and g). Together these results indicate that the epigenomic status of genes can be used to identify critical genes necessary for multicellularity. Interestingly histone deacetylases of the *hdaD* family have been shown to bind to MADS box transcription factors on DNA (*39, 40*), raising the possibility of a coordinated evolution of transcription factor networks and epigenetic machinery for regulating multicellularity.

## Discussion

### *Dictyostelium*: a new epigenetic model organism for studying the evolution of multicellularity

By comparing the chromatin accessibility at syntenic genes of *Dictyostelium* to *S. pombe* and *C. elegans*, we were able to identify a set of candidate genes whose chromatin status correlated with unicellularity or multicellularity. This analysis identified one gene, *srfA,* which had previously been shown to play a role in developmental timing (*24, 25*), as well as several uncharacterized genes. We found that knocking out three novel “multicellular” genes delayed or inhibited multicellularity without making the *Dictyostelium* less fit. Similarly, the overexpression of one of these “multicellular” genes was sufficient to triple the number of cells integrated into a multicellular state. Together the approaches employed in this study have identified several genes, based on their epigenetic status, which are necessary and sufficient for the transition to multicellularity. This analysis also suggests that a deeper comparison of not only the chromatin accessibility at promoters, as was performed in this study, but also the chromatin accessibility in the gene body or further upstream of the TSS along with specific chromatin modifications across different unicellular and multicellular organisms and *Dictyostelium* stages could reveal additional genes essential for multicellularity. It will be important, in future experiments, to also examine whether changing the epigenome at these critical genes is sufficient to induce multicellularity.

Since *Dictyostelium* rapidly transitions between physiologically and morphologically distinct states, this organism is ideally situated to be utilized as an epigenetic model organism for examining some of the critical outstanding questions in the field of epigenetics. Here we have displayed the ease and capacity to not only perform comparative observational science but also to manipulate the genome and epigenome of *Dictyostelium* to test theories. Therefore, we feel this will be a useful organism to study a host of epigenetic questions beyond the role of epigenetics in the evolution of multicellularity. Our study has provided a comprehensive epigenomic sequencing of critical stages of *Dictyostelium.* It also demonstrates how this comprehensive dataset can be used to study the role of epigenetics in multicellularity. This comprehensive dataset opens the possibility of identifying additional processes that epigenetic modifications and drugs can regulate.

### Proof-of-principle that epigenetics plays a role in regulating multicellularity (Genetic vs epigenetic)

In this study, we used *Dictyostelium discoiduem* as a model organism to investigate the role of epigenetics in the evolution of multicellularity. Through a combination of chemical, comparative genomic, and epigenomic approaches, we identified several key components of multicellular development. The histone deacetylase inhibitor, TSA, had previously been demonstrated to delay development in *Dictyostelium* (*25*), suggesting that histone acetylation could play a critical role in development. We confirmed these findings and found that chemical inhibitors of histone methylation, Sinefungin and BIX 01294, also delayed development, suggesting that chromatin regulation is important for multicellularity. We also examined the distribution of several histone modifications; H3K4me3, H3K4me1, H3K27ac, H3K36me3, H3K27me3 and H3K9me3. Principal component analysis (PCA) demonstrated that specific histone modifications, such as H3K4me3 and H3K27ac, were sufficient to distinguish unicellular from multicellular forms of *Dictyostelium*. The importance of H3K4me3 in regulating multicellularity in *Dictyostelium* is reinforced by the demonstration that deletion of the H3K4me3 methyltransferase *set1*, causes an unusually rapid development (*18*). Our results further demonstrated the importance of H3K27ac by our unbiased identification that the histone deacetylase *hdaD,* which we demonstrate is an H3K27 deacetylase (Fig. 5, g-i), increases expression upon transition to multicellularity (Fig. 2d) and is necessary for an appropriately timed induction of multicellularity (Fig. 4, a and b). Together these results suggest that chromatin modifications are important and could help to drive multicellularity in *Dictyostelium*.

To gain a global view of chromatin accessibility of *Dictyostelium,* we performed ATAC-seq at specific life cycle stages. We found dramatic differences in chromatin accessibility in the unicellular versus multicellular stages, suggesting that chromatin reorganization during this transition to multicellularity is critical. This analysis revealed that MADS-box transcription factor motifs become accessible upon transition to multicellularity (Extended Data Fig. 4). The importance of these binding sites becoming accessible is reinforced by previously published work demonstrating that two MADS-box transcription factors, *mef2A* and *srfA,* are necessary for appropriate multicellularity timing (*24–26*). These results suggest that to transition to multicellularity, *Dictyostelium* reorganizes its chromatin structure to make binding sites for multicellular specific transcription factors *mef2A* and *srfA* accessible for binding, to facilitate the transcription of their target genes. Excitingly, class II histone deacetylases, of which *hdaD* is a member, have been shown to bind to MADS box transcription factors on DNA (*39, 40*), raising the possibility of a coordinated evolution of transcription factor networks and epigenetic machinery for regulating multicellularity. While our work, and the work of many other labs, has demonstrated that genetic changes helped contribute to the evolution of multicellularity, this study adds to this work and suggests that these genetic changes were accompanied by contemporaneous epigenetic changes and suggests that the co-evolution of new transcription factor networks and chromatin remodeling coalesced to facilitate multicellularity.

## Acknowledgments

We thank Peter Devreotes, Marc Edwards, Budri Sharif and other members of the Devreotes lab for the *Dictyostelium* strain and experimental advice about *Dictystelium* culturing conditions. We thank J. Lieberman for critical reading of the manuscript.

## Funding

S.Y.W. was supported by a Croucher Foundation fellowship. E.A.P. was supported by an LSRF fellowship. K.T. was supported by a JSPS fellowship. P.L.G. was supported by grants from the Searle Scholars Program, Rita Allen Foundation, and by an NIH grant (DP2AG067490). This research was supported by NIH grants (R00AG043550 and DP2AG055947) to E.L.G.

## Author contributions

S.Y.W., E.A.P., and E.L.G. conceived and planned the study and wrote the paper. All analysis and chemical and genetic manipulation of *Dictyostelium* experiments were performed by S.Y.W.. E.A.P. performed ChIPseq and ATACseq experiments. I-H.W. performed single cell RNAseq analysis. H.B. helped with chemical manipulation of *Dictyostelium* and purification of HdaD experiments. K.T. performed apoptosis experiments. C.F. optimized chemotaxis assays and helped produce Extended Data Fig. 8. G.S. helped produce Figs. 4d and Extended Data Fig 1. P.L.G. advised I-H.W. and helped write the paper. E.L.G. generated staged *Dictyostelium* for RNAseq, ChIPseq, and ATACseq experiments and performed chromatin immunoprecipitations. All authors discussed the results and commented on the manuscript.

## Competing interests

Authors declare no competing interests.

## Data and materials availability

All ChIPseq, RNAseq, and ATACseq datasets are available in the GEO database accession # GSE137604.

Unprocessed and uncompressed imaging data is available at https://data.mendeley.com/datasets/zfhh6xzck4/draft?a=43501cd3-097c-4456-b8a7-ba08a4c758b7.

## Methods

### Strain

The Axenic strain (AX2) of *Dictyostelium discoideum* was generously given by Peter Devreotes at Johns Hopkins University. This stain was used as the main material for RNA-seq, ChIP-seq and ATAC-seq experiments. The other axenic strain (AX4) and the *srfA* knock out mutant was obtained from Dicty Stock Center. We cultured the strain at 22 °C shaken in the incubator at 180 rpm in HL5 medium with 300 μg/ml streptomycin.

### Antibodies

Antibodies used for histone chromatin immunoprecipitation and western blots were: rabbit anti-H3 (Abcam Ab1791), rabbit anti-H3K27ac (Abcam Ab4729), rabbit anti-H3K4me3 (Abcam Ab8580), rabbit anti-H3K9me3 (Abcam Ab8898), rabbit anti-H3K27me3 (Abcam Ab6002), rabbit anti-H3K36me3 (Abcam Ab 9050), rabbit anti-H3K4me1 (Abcam Ab8895). These antibodies have demonstrated specificity in eukaryotes (*41–47*).

### RNA-seq

mRNA was extracted from AX2 cells stored at −80 °C using Dynabeads mRNA direct micro purification kit (Invitrogen). Samples were lysed using tissue homogenizer (Kontes) in the lysis buffer. Two biological replicates were assigned for each vegetative, streaming, mound, and fruiting stage respectively. In total 8 groups were prepared for RNA-seq libraries with 500 ng mRNA as starting materials using NEXTflex Illumina qRNA-Seq Library Prep Kit (Bioo Scientific). In short, mRNA was fragmented in a cationic buffer and underwent first and second strand synthesis, adenylation, molecular indexed adapter ligation and PCR amplification for 11 cycles. The amplified products were cleaned up by Agencourt AmPure XP Magnetic Beads (Beckman Coulter) and quality checked by Agilent 2200 Tapestation D1000. The optimal cluster density was determined by KAPA library quantification kit before pooling the samples into a pooled library (5 nM) and sequencing through Nextseq 500 platform.

The fastq files were filtered with Q score 30 or above and were trimmed for the sequencing adaptors using CutAdapt. The analysis pipeline started with pseudo-alignment of the reads using kallisto(*48*), followed by building an index and quantification process, which generated h5 files containing the raw binary files for preliminary differential analysis in sleuth (*49*). Differential analysis were conducted in EdgeR *(50)*, briefly, the DEGlist object were created, and genes were kept that achieved at least one count per million (cpm) in two biological replicate samples. The effective library size were computed using Trimmed Mean of M values (TMM) normalization. The significant genes were deducted by computing NB exact genewise tests with corrected *p* < 0.05. The gene ontology (GO term) for over-representation analysis and diagrams were generated by R package clusterprofiler (*51*). Sequencing QC matrix is shown in Table S8.

### ChIP-seq

10 million *D. discoideum* cells were harvested and dissociated at different life cycle stages as described in (*52*) with the following modifications. After dissociation cells were cross-linked with 1x linking buffer (11 mM HEPES-NaOH, 110 mM NaCl, 1.1 mM EDTA and 1.1 mM EGTA) with 1% formaldehyde for 10 minutes at RT and quenched by 2M glycine. The samples were fragmented by applying a biorupter for 25 cycles in the cold room with 30 seconds on and 30 seconds off per cycle. The precleared chromatin was incubated with antibody-coupled bead slurries and rotated at 4°C overnight. DNA was eluted from the beads by 200 μl TE with 1% SDS.

Histone ChIP-Seq libraries were generated using the Ovation Ultralow V2 kit (Nugen, 0344-32) according to the manufacturer’s instructions. Libraries were PCR amplified for 13-16 cycles depending on the antibody, and library quality was assessed using the Agilent 2100 Bioanalyzer (Agilent Technologies). Seventy-five bp reads were generated on the Illumina Nextseq 500 (Illumina). ChIP-seq were sequenced to a depth of at least 8 million total reads/sample.

The raw fastq files were trimmed for the first and last 8 nucleotides, the multiqc was use to summarize the fastQC results (*53*), and the trimmed files were mapped to the *Dictyostelium discoideum* genome (*12*) using bowtie2 (*54*). The duplicated reads were removed by picard and sorted bam files were subject to peak calling with MACS2 (*55, 56*). Both DNA input and H3 were used as controls. The peaks were visualized by Integrative Genomics Viewer (IGV 2.3.44). Differential binding analyses were performed by R package Diffbind using log2 normalized ChIP read counts with FDR < 0.05 (*57*). The metagene analysis and genomic feature distribution were conducted by ChIPSeeker using anotatePeak function with narrowpeak files as input (*58*). Transcription start site (TSS), exon, intron and transcription termination site (TTS) were determined by using makeTxDbFromBiomart function from the Ensemble BioMart database (version 2.7). Promoters were defined by the regions 1500 bp upstream to the TSS, and downstream was defined as the 3000 bp downstream of gene end (TTS). Since some annotation may overlap, we adopt the default principle in ChIPSeeker to prioritize genomic annotation in the following order: promoter, exon, intron, downstream and intergenic. Sequencing QC matrix is shown in Table S8.

### ATAC-seq

To assess regions of open chromatin, frozen pellets of *D discoideum* cells from the various stages were washed twice with cold PBS and lysed with 10 mM Tris-HCl (pH 7.4), 10 mM NaCl, 3 mM MgCl_2_, 0.1% Igepal-630. Nuclei were transposed using the Nextera DNA Library Prep Kit (Illumina, FC-121-1030). Transposed libraries were amplified for 11 cycles and size selected for fragments ranging 200-1000 bp. ATAC-seq experiments were sequenced to a minimum depth of 10 million reads/sample. For preparation of fresh samples, *Dictyostelium* were physically dissociated and washed twice with PBS with 0.04% BSA and lysed with 10 mM Tris-HCL (pH 7.4), 10 mM NaCl, 3 mM MgCl_2_, 0.5% Tween 20, 0.5% NP-50, 0.05% digitonin (Promega G944A), 1% BSA, and 0.5% cellulase on ice or room temperature for 30 minutes to fully lyse the cells. Validation of equivalent ATACseq lysine of different stages was performed by trypan blue staining and optimized by real-time PCR detection of ribosomal RNA release.

The raw fastq files were trimmed for the first and last 8 nucleotides, the multiqc was use to summarize the fastQC results (*53*), and the trimmed files were mapped to the *Dictyostelium discoideum* genome (*12*) using bowtie2 (*54*), and the mitochondria genome was removed in the alignment process. The duplicated reads were removed by picard and sorted bam files were subject to peak calling with MACS2 (*55, 56*). The peaks were visualized in Integrative Genomics Viewer (IGV 2.3.44). Likewise, open chromatin regions and peak annotation were deducted with Diffbind (*57*) and ChIPSeeker, respectively (*58*). Sequencing QC matrix is shown in Table S8.

### Motif analysis

The stage/unicellular/multicellular specific open chromatin regions, enriched histone binding sites, and the promoters of significant enriched genes were extracted as the input sequence for homer (*59*). All the promoter sequences were obtained from dictyBase as the background sequence (*60*). The motif discovery was performed using “findMotif.pl” command from homer.

### Syntenic analysis

The phylogenetic tree was constructed on interactive tree of life (iTOL)(*61*). The syntenic regions/genes among *D. discoideum*, *C.elegans* and *S.pombe* were obtained using metaphors based on phylogeny orthology prediction at genomic scale (*34*). The comparison among the species and intersection regions were extracted using bedtools (*62*). The graphs were generated on R, excel or prism.

### Developmental Assay

The *D. discoideum* cells were harvested in mid-log phase for 5*10^8^ cells. The cells were suspended in developmental buffer (5 mM Na_2_HPO_4_, 5 mM KH_2_PO_4_, 1 mM CaCl_2_ and 2 mM MgCl_2_) and washed 3 times. The developmental assays were performed on KK2 agar plates. 100 nM, 500 nM and 1000 nM chemical inhibitors (Trichostatin A, T8552; BIX 01294 trihydrochloride hydrate, B9311; (R)-PFI-2, SML1408p; Sinefungin, ab144518) were added to the KK2 agar plates. The picture capture time points for the development assay with chemical treatment were set at 8 hours, 16 hours, 19 hours, and 24 hours. The picture capture time points for the development assay with mutant strains were set at 8 hours, 16 hours, 21 hours, 25 hours, and 41 hours.

### Cell number quantification and morphological measurement

To quantify the germ cell number in the fruiting bodies, the assay was performed as in (*63*). Germ cell number from individual fruiting bodies were collected by pipette and were placed in 20 μl PBS. The cells were counted with a hemocytometer (Hausser Scientific Cat.#3520). The diameter of individual mound and fruiting bodies were measured using stereo Discovery V8 microscope.

### Chemotaxis Assay

Chemotaxis assays were performed as in (*64*). Briefly the protocol was modified as follows. The *D. discoideum* cells were cultured in a density of 2×10^6^ cells/ml for the purpose of the experiment. The cells were pelleted in a 1X KK2 buffer (2.2 g KH_2_PO_4_, 0.7 g K_2_HPO_4_). The cells were starved for 5 – 8 hours in a shaker and resuspended at a concentration of 1×10^8^ cells/ml. For making the chemotaxis plates, 4 wells were made in the 1X KK2 agar plates (1X KK2 buffer and 1.5g Agar). 4 small dots at 6 mm from the edge of each well were marked on the agar plates for showing the location of cells deposition. 20 μl of 250 μM of folic acid (Millipore Sigma CAS 59-30-3) dissolved in 100 mM NaOH were injected into the wells by a P10 pipette for 20 minutes absorbing by the agar. 1 μl of *D. discoideum* were loaded on the dots and the plates were incubated at room temperature in a humidified chamber. The movement of cells were captured after 5 hours using Zeiss steoREO Discovery V8 microscope. The number of cells were counted in each direction (north, south, east and west) in a 30 degree angle range and plotted in Prism 8 (Version 8.1.1).

### Annexin V, Propidium iodide, and Trypan blue staining

Cell viability was examined by trypan blue (VWR), propidium iodide (BD Biosciences), and Annexin V staining (BD Biosciences). For Annexin V staining, Cells were washed twice with cold PBS and suspended in 1X Binding Buffer (10 mM Hepes pH 7.4, 140 mM NaCl, and 2.5 mM CaCl_2_) at a concentration of 1 million cells/ml. 100 μl of the solution were transferred to a new tube and 5 μl of FITC Annexin V was added to the cells and incubated for 15 minutes at room temperature in the dark. The samples were imaged using a fluorescent microscope (Nikon Eclipse E600) with 485/535 nm (FITC) filter and images were analyzed by ImageJ (1.0).

For trypan blue staining, 10 μl 0.4% solution of trypan blue was mixed with 10 μl of cells at 5E5 cells/ml. The mixture was counted on a hemacytometer immediately after mixture.

For PI staining, 1E6 *D. discoideum* cells were treated with 1 μM TSA, 5 μM sinefungin, or 10 μM BIX 01294 for 6 hours. Cells were subsequently washed and resuspended with cold PBS supplemented with 5% FBS and stained with 2 mg/ml Propidium Iodide (BD Biosciences) at room temperature for 10 min. Stained cells were fixed with 1% PFA. Samples were analyzed by BD LSR Fortessa (BD Biosciences) and data analysis was performed using Flow Jo (Tree Star).

### CRISPR-Cas9 knock-out experiment

The all-in-one sgRNA and Cas9 expression vector pTM1285 were obtained from Quantitative Biology Center (QBIC) Riken, Japan. The sgRNAs were designed on CRISPOR tool(*65*). sgRNA were ligated to predigested pTM1285 with BbsI (NEB), and stable competent cells were transformed with the resulting construct(NEB, C3040). Ligation was confirmed by PCR and Sanger sequencing. Primers used were sense oligo for the target 5′-AGCAN(1)NNNNNNNNNNNNNNNNNNN(20)-3′ and tracrRNA 5′-AAGCTTAAAAAAAGCACCGACTCGGTGCC-3′. The sequencing primer was 5′-AAGCTTAAAAAAAGCACCGACTCGGTGCC-3′. The validated plasmids were transformed into the cells using calcium phosphate precipitation method (*66*). Briefly, 10 μg plasmid DNA is co-precipitated with 60 μl of 1.25 M CaCl_2_ in HBS buffer (4.0 g NaCl, 0.18 g KCl, 0.05g Na_2_HPO_4_, 2.5 g HEPES and 0.5 g D-glucose, pH adjusted to 7.1). The DNA solution was added to the plates for 30 minutes and then 12.5 ml Bis-Tris (2.1 g Bis-Tris, 10 g proteose peptone, 5 g yeast extract, and 10 g D-glucose in distilled water, pH adjusted to 7.1 with HCl) for 4 – 8 hours. The condensed DNA on cell surface was transformed into the cells by 15-18% glycerol for exact 5 minutes and incubated with 12.5 ml Bis-Tris solution overnight to allow the expression of the selection marker. The medium was replaced by HL5 with 10-15 μg/ml of G418 after 16 hours, and cultured for another 3 days. The cells were plated on SM agar paltes with *K. aerogenes* and incubated for 3 – 4 days until plaque formed. The cells were transferred and expanded in the HL5 medium with 300 μg/ml streptomycin. Knock-out strains were validated by sanger sequencing.

### Overexpression *Dictyostelium* strains

cDNA of *hdaD* and *smcl1* was cloned into pJSK123 over-expression plasmid (*67*) between BamHI and AvrII sites. The recombinant plasmids were transformed into cells using calcium phosphate precipitation method (*66*). Briefly, 10 μg plasmid DNA is co-precipitated with 60 μl of 1.25 M CaCl_2_ in HBS buffer (4.0 g NaCl, 0.18 g KCl, 0.05g Na_2_HPO_4_, 2.5 g HEPES and 0.5 g D-glucose, pH adjusted to 7.1). The DNA solution was added to the plates for 30 minutes and then 12.5 ml Bis-Tris (2.1 g Bis-Tris, 10 g proteose peptone, 5 g yeast extract, and 10 g D-glucose in distilled water, pH adjusted to 7.1 with HCl) was added for 4 – 8 hours. The condensed DNA on the cell surface was transformed into the cells by 15-18% glycerol for 5 minutes and incubated with 12.5 ml Bis-Tris solution overnight to allow the expression of the selection marker. The medium was replaced with HL5 with 10-15 μg/ml of G418 after 16 hours, and cultured for another 3 days. The cells were then plated on SM agar plates with *K. aerogenes* and incubated for 3 – 4 days until plaques formed. The cells were transferred and expanded in the HL5 medium with 300 μg/ml streptomycin. Over-expression strains were validated by real-time PCR.

### Quantitative real-time PCR(qRT-PCR)

The differential expression of candidate genes were analyzed using qRT-PCR. Total RNA was extracted from *D. discoideum* cells using Invitrogen PureLink RNA (Invitrogen, 12183025). First strand cDNA synthesis was conducted using SuperScript III First-Strand Synthesis (Invitrogen, 18080051). qRT-PCR of cDNA was performed using iTaq Universal SYBR Green Supermix (Bio-Rad, 172-5122) on a CFX96 Real-Time System (Bio-Rad). The PCR assay contained 1:4 diluted reverse transcription products, 1X master mix, and 200 nM of each primer, and was performed with 5-min initial denaturation at 95°C followed by 40 cyles of 95°C for 5s and 60°C for 45s. The mtRNA of *D. discoideum* was used for internal normalization for each gene. The comparative Ct method (ΔΔCt) was used to calculate gene expression. The primer sequences of the selected genes were designed using Macvector (15.5.0). The following gene-specific primers were used: mtRNA-forward, 5’-gggtagtttgactggggcgg-3’, mtRNA-reverse, 5’-cactttaatgggtgaacacc-3’; DDB_G0283437-forward, 5’-gagggtgtataggtacatttgcagag-3’, DDB_G0283437-reverse, 5’-tgggttccaatttcccaatcccaaac-3’; DDB_G02813787-forward, 5’-tccctaacaccaacatcaaccaacga-3’, DDB_G02813787-reverse, 5’-tatacatgaccagtttcggaggcaac-3’; DDB_G0288013-forward, 5’-acaagaaatcgattgggagatgt-3’, DDB_G0288013-reverse, 5’-tggcatttgttcttggatttttct-3’; DDB_G0279267-forward, 5’-tggctcaccaataccatcacctccat-3’, DDB_G0279267-reverse, 5’-cgagttcgaactattgctgttg-3’; DDB_G0274899-forward, 5’-ccagagggaaataatagtcctggt-3’, DDB_G0274899-reverse, 5’-ctccaattcgaaccacaaac-3’; DDB_G0267824-forward, 5’-gcagctggtagtagttcatcca-3’, DDB_G0267824-reverse, 5’-tgtgcatattcttctttaccctcaa-3’; DDB_G0277333-forward, 5’-tggcatttctcaggcagtttcgacca-3’, DDB_G0277333-reverse, 5’-ccaacatcagtgagggtattca-3’; DDB_G0293674-forward, 5’-accaacgtgctcttcgtcgtttgag-3’, DDB_G0293674-reverse, 5’-ggaacgatgtttgtcattacaccac-3’; DDB_G0294934-forward, 5’-gggtagtttgactggggcgg-3’, DDB_G0294934-reverse, 5’-cactttaatgggtgaacacc-3’; DDB_G0279267_OE-forward, 5’-tatggatccatgtcaacaattcatc-3’, DDB_G0279267_OE-reverse, 5’-tatcctaggttaatcgtcttcgctta-3’; DDB_ G0288013_OE-forward, 5’-tatggatccatggatattgcacaatcaattca-3’, DDB_ G0288013_OE-reverse, 5’-tatcctaggctataaagtttttaagaaattttg-3’, DDB_G0279267_Cat-forward, 5’-caacaatcatcacaattacc-3’, DDB_G0279267_Cat-reverse, 5’-tatcctaggttaatcgtcttcgctta-3’.

### Western blot analysis

Cells were lysed with RIPA buffer and quantified by Bradford assay (Biorad 5000006). 5 to 10 μg of proteins were loaded for gel electrophoresis, and transferred to nitrocellulose membrane (Bio rad 1620097) with 100V for 1.5 hours. The membranes were incubated in 5% milk for 1 hour at room temperature and incubated with primary antibodies (1:1000 dilution). After washing with TBST 3 times, each for 10 minutes, the membranes were incubated with secondary antibody (Millipore 3136001, Rabbit IgG, 1:3000 dilution). The blots were reacted with chemiluminescent HRP substrate (Millipore WBKLS0500) and imaged in ChemiDoc Touch Imaging System (Bio rad).

### Deacetylation assays

The coding sequence of the catalytic domain of *hdaD* was cloned as an in-frame fusion to the His tagged vector pET28a. The recombinant protein was expressed in *E. coli* BL21. Overnight induction of protein expression was carried out with 1 mM IPTG at 18°C. Bacteria were harvested at 4000 rpm, 4°C and resuspended in 10 mL protein purification lysis buffer (50 mM Tris-HCl pH 7.5, 0.25 M NaCl, 0.1% Triton-X, 1 mM PMSF, 1 mM DTT, 20 mM imidazole, and protease inhibitors). After freezing the pellet at −80°C for 1 hr, the lysate was sonicated with a Bioruptor for 5 mins on high level with 30 seconds on and 30 seconds off. Proteins were purified with Ni-NTA beads (Qiagen). Proteins and beads were washed 3 times with protein purification lysis buffer before incubating the beads with elution buffer (400 mM imidazole in protein purification lysis buffer, pH 8.0) for 30 mins. Eluates were dialyzed overnight at 4°C with dialysis buffer (50 mM Tris-HCl pH 8.0, 1mM EDTA, 1mM DTT, and 20% glycerol). Bradford assays and SDS-page gel electrophoresis followed by coomassie staining was performed to determine integrity and quantity of purified proteins. Histone proteins were purified from *D. discoideum* using the EZExtract core histone isolation kit (BioVision). *In vitro* deacetylation reactions were performed with 10 μg of histone proteins and 5 μg of *hdaD* protein or 5 μg GST-RPA-2 protein as a control. The deacetylation reaction was performed in a buffer solution (50 mM Tris-HCl, 4mM KCl, 4 mM MgCl2, 0.2 mM DTT, 1 mM NAD^+^) and incubated at 30 °C for 30 minutes. Western blot analysis was performed as previously described.

### Single Cell RNA sequencing data analysis

*D. discoideum* was dissociated into single cells from different stages by physical pipetting with PBS + 0.04% BSA. For each sample, the cells were counted and diluted to 1000 cell/μl in 1X PBS with 0.04% Bovine Serum Albumin (BSA) prior loading to 10X Genomics Chromium machine. 10X Chromium Single Cell 3′ v3 reagent kits were used according to the manufacturer’s instructions to generate scRNAseq libraries. Libraries were sequenced paired-end 150bp on Illumina Hiseq4000/Novoseq (Novogene).

### Single Cell RNA sequencing data analysis

Fastq files were processed with Cell Ranger (version 3.1.0) (https://support.10xgenomics.com) on the DolphinNext pipeline builder (*68*) to generated the merged gene-barcode matrix. Matrix was loaded into R (version 3.6.3) and analyzed mainly with Seurat package ((*69*), version 3.1.5). Low quality cells or outliers were filtered out from the data by the following threshold: 200 < total Gene count < 7000, 500 <total UMI count <30000. Filtered matrix were normalized with SCTransform (*70*) and reduced the dimension with principal component analysis. The batch effect were corrected by harmony (*71*).The significant principal components (PCs) are used to generate the UMAP plot for visualization. Clustering were perform by Louvain algorithm based on K-nearest neighbor graph. Hurdle model tailored was used to perform differential expression analysis of the individual clusters (*72*). Gene Ontology (GO) analysis was performed with clusterProfiler (*51*).

For trajectory inference analysis, the cells from vegetative, slug and fruiting stage were down sampled and merged. The uninformative genes were filtered out by keeping the genes expressing 3 UMIs per cell for at least 10 cells. The filtered matrix were normalized with full quantile normalization and reduced the dimension with PCA. The first 3 PCs were used for k-means clustering and subsequent analysis. Slingshot were performed to predict the development lineage (*73*). The generalized additive model were used to identify the temporally expressed gene through the pseudotime.

## Notes

### Competing Interest Statement

The authors have declared no competing interest.

