## Supplementary Figures 1-13 for "Role of Epigenetics in Unicellular to Multicellular Transition in *Dictyostelium*"

### Supplementary Information

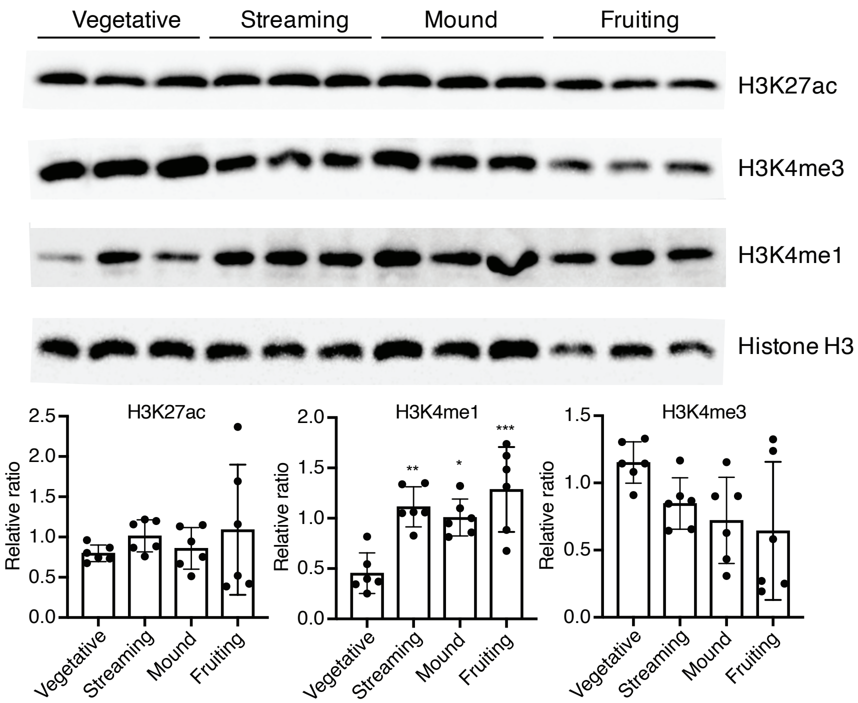


**Extended Data Fig. 1 Chromatin modifications change during *D. discoideum* development**
A decrease in H3K4me3, no change in H3K27ac, and an increase in H3K4me1 is observed by western blot analysis of three biological replicates of *D. discoideum* analyzed at four different developmental time points. Quantification of two independent experiments performed in triplicate is depicted below. *: $p<0.05$, **: $p<0.005$, ***: $p<0.001$, as assessed by multiple comparison one-way ANOVA analysis.

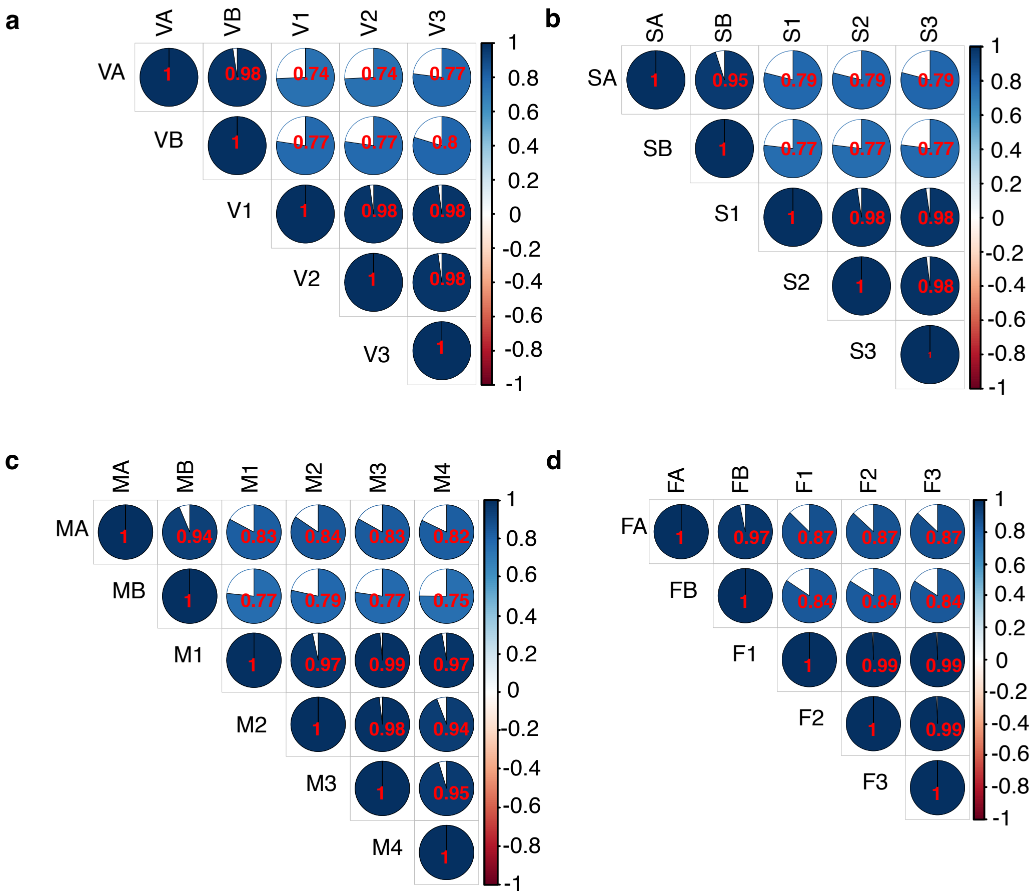


**Extended Data Fig. 2 ATACseq analysis displays a high degree of reproducibility between replicates of a, vegetative, b, streaming, c, mound, and d, fruiting body stages.**

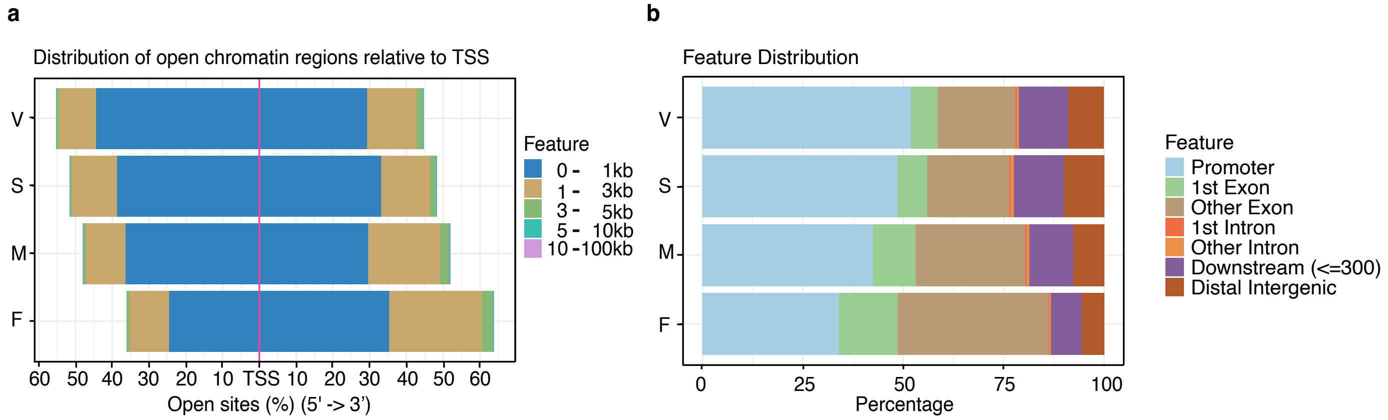


**Extended Data Fig. 3 ATACseq analysis of changes in accessible chromatin at different life cycle stages**

**a**, An analysis of chromatin accessibility in relation the nearest TSS demonstrates that unicellular *D. discoideum* have more accessible chromatin further away from the TSS than do multicellular stages. **b**, An analysis of the chromatin accessibility at specific genomic features at different developmental stages reveals that the unicellular stage has a lower degree of accessibility in the promoter and a higher degree of accessibility in exons than multicellular stages.

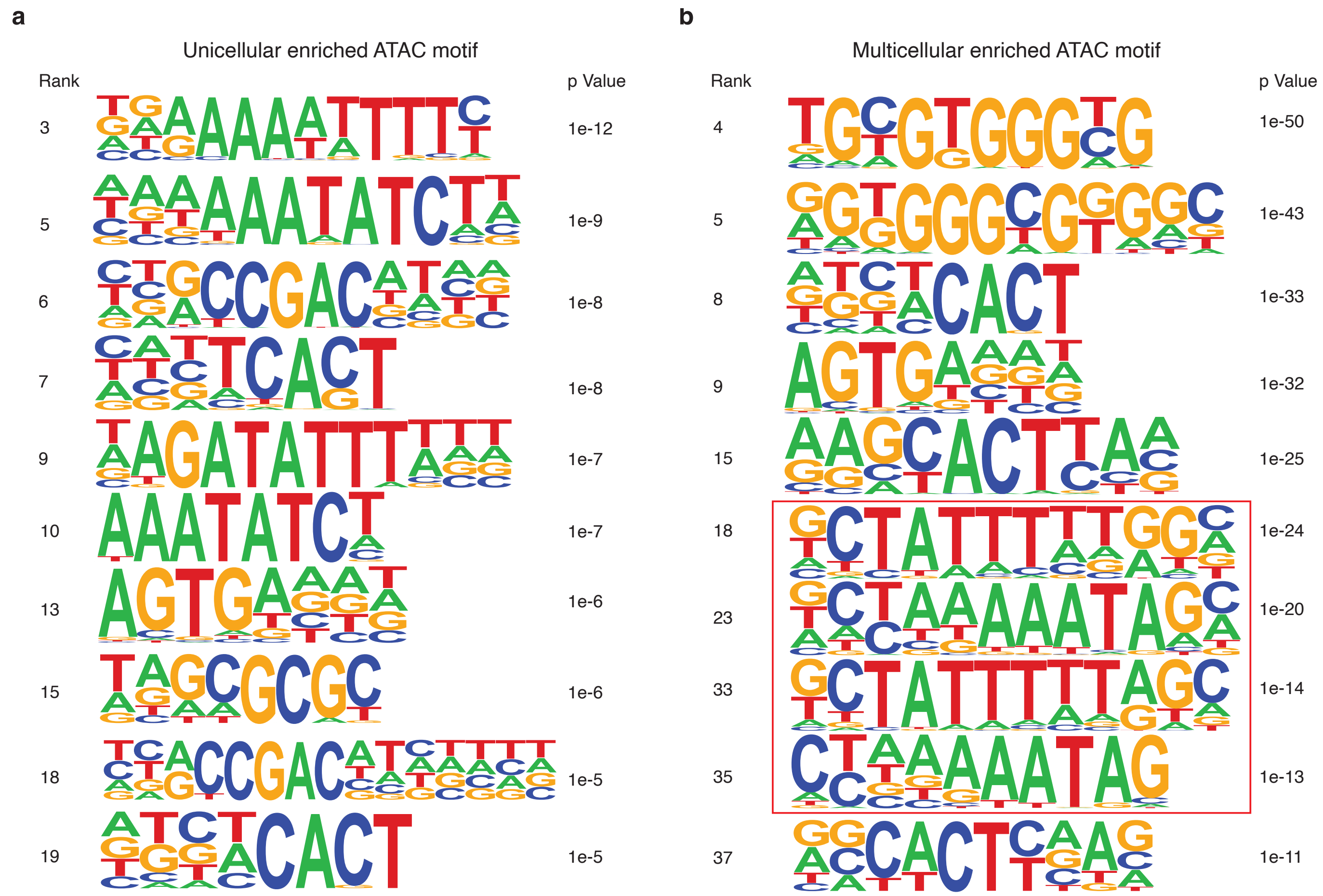


**Extended Data Fig. 4 a, Unicellular and b, multicellular stages display unique enriched DNA sequence motifs during ATACseq analysis**

Comprehensive ATACseq motif analysis can be accessed at https://www.dropbox.com/sh/flf9wqvzz4go7dc/AADr3KXeZQoIr9JjWgyhcikoa?dl=0.

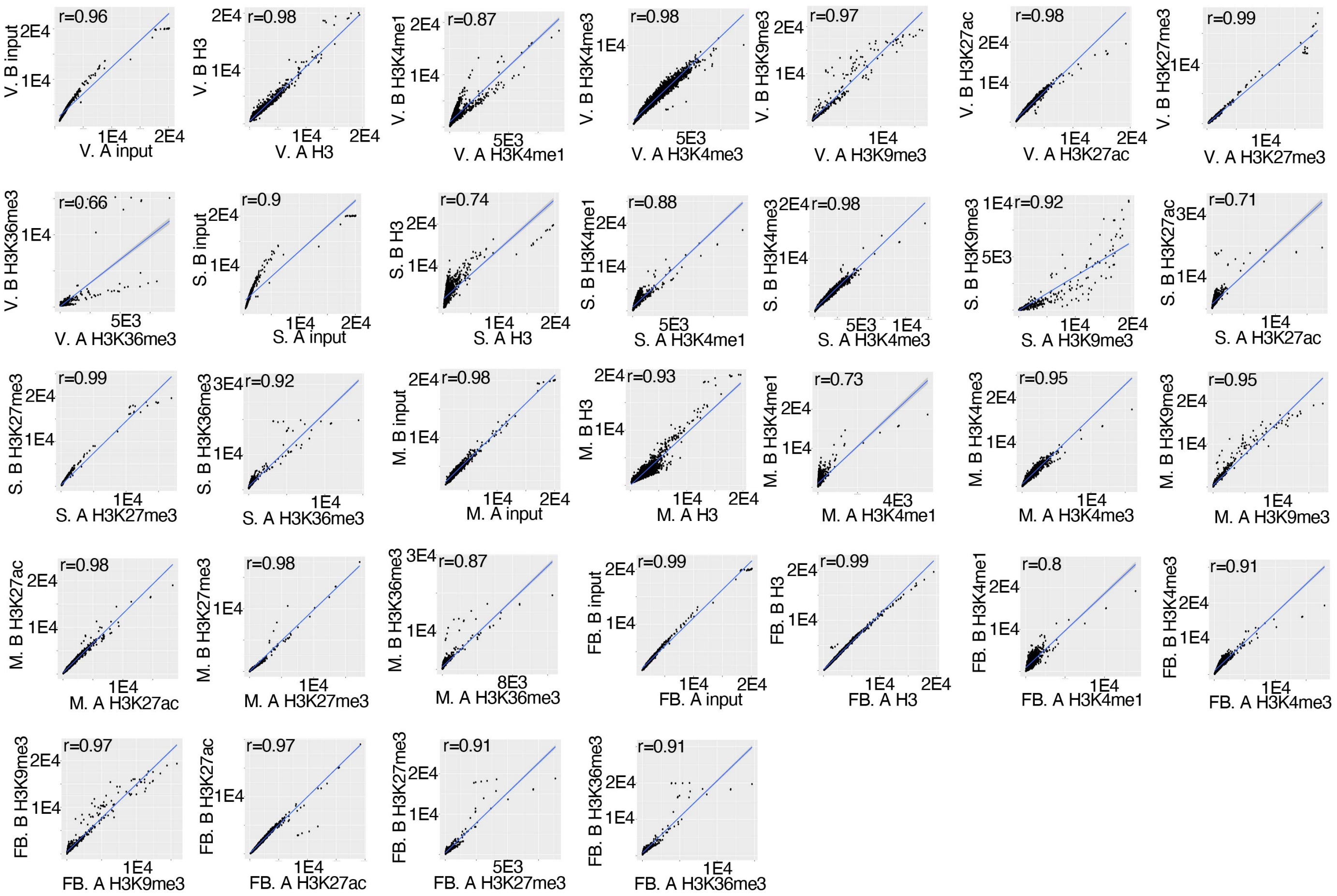


**Extended Data Fig. 5 ChIPseq analysis displays a high degree of reproducibility between replicates of 7 different chromatin immunoprecipitations for vegetative, streaming, mound, and fruiting body stages.**

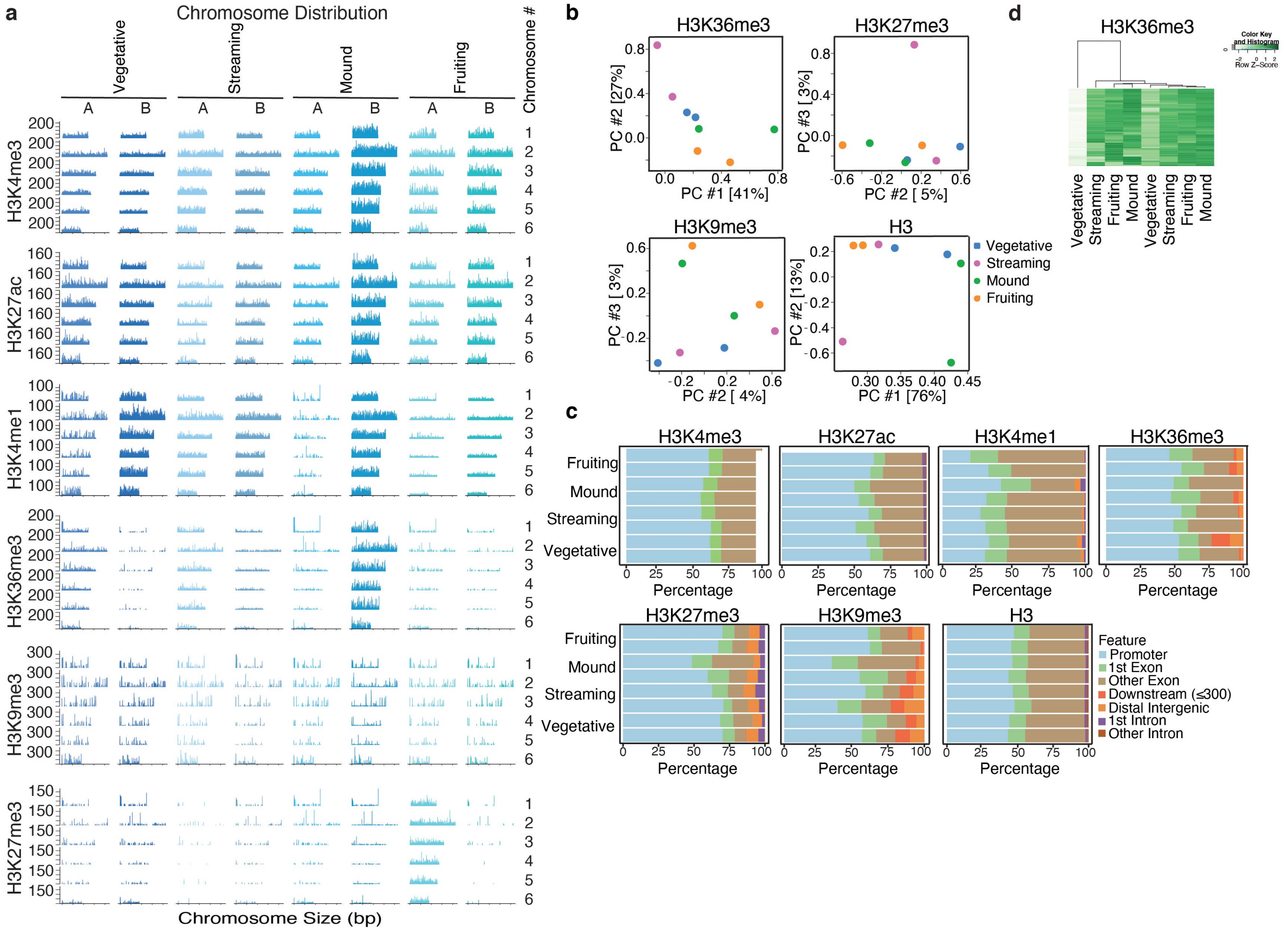


**Extended Data Fig. 6 Chromosome distribution, Principal Component analysis, and Feature Distribution of ChIPseq data for 6 chromatin modifications.**

**a,** Chromosome distribution of ChIPseq data for 6 chromatin modifications are displayed across all 6 chromosomes. **b**, PCA of ChIPseq analysis of H3K27me3, H3K9me3, and H3 reveals that these modifications are insufficient to distinguish unicellular from multicellular *D. discoideum*. **c**, An analysis of chromatin modifications distribution at genomic features in the *D. discoideum* genome reveals that chromatin modifications display differences in feature distribution. **d**, H3K36me3 heatmaps did not display distinct patterning in unicellular (vegetative) and multicellular (mound and fruiting body) stages.

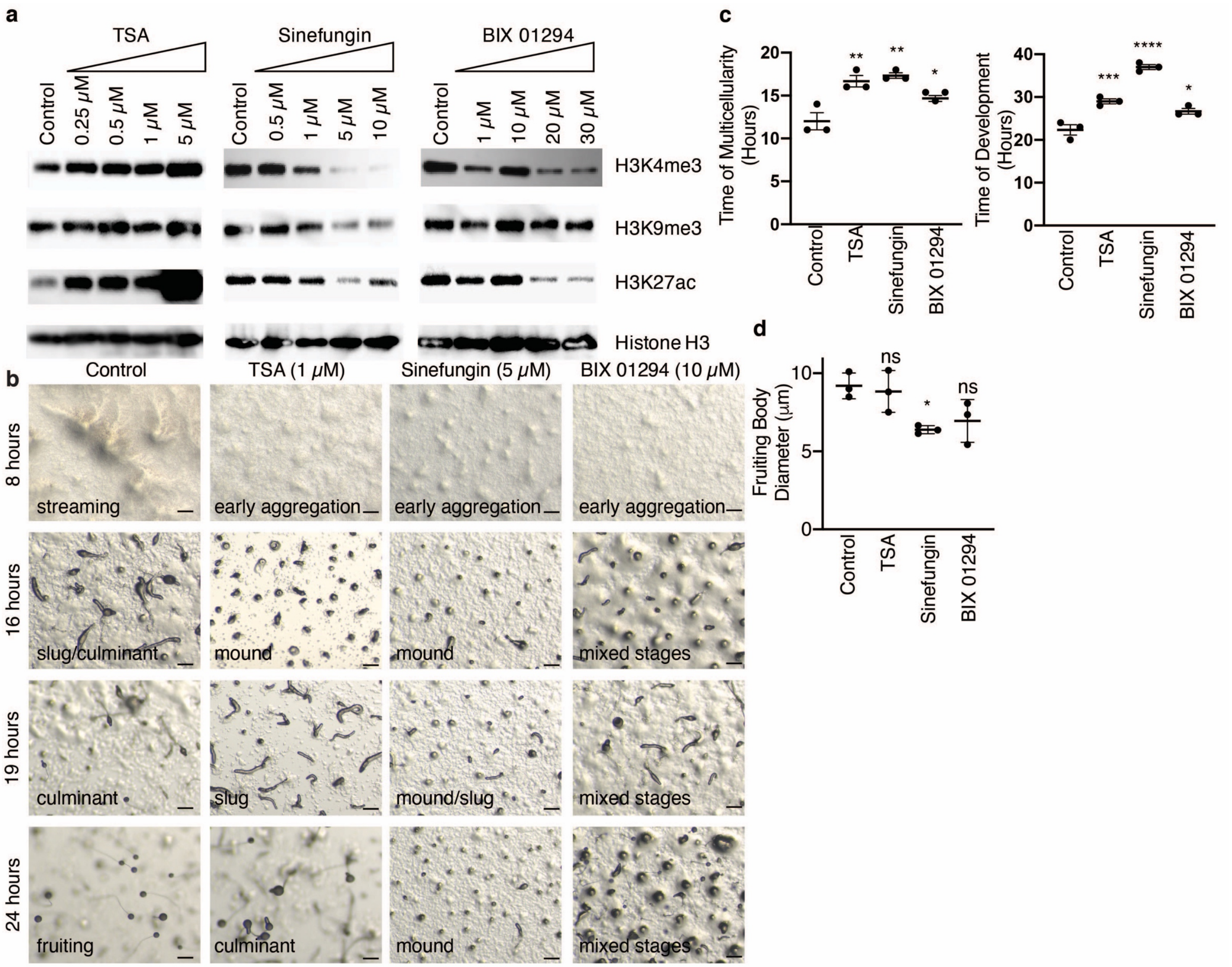


**Extended Data Fig. 7. Chemical inhibitors of chromatin modifications delay *D. discoideum* development**

**a**, TSA, Sinefungin, and BIX 01294 each display a concentration dependent inhibition of H3K27ac, H3K4me3, H3K9me3 and H3K27ac, and H3K4me3 and H3K27ac respectively as assessed by western blotting. **b**, Representative pictures of *D. discoideum* development on KK2 plates show developmental delay after treatment with 5 μM sinefungin, 10 μM BIX 01294, and 1 μM TSA at 8, 16, 19, and 24 hours. Stage is displayed in the lower right hand corner. Scale bars are 20 microns. **c**, Treatment with 5 μM sinefungin, 10 μM BIX 01294, and 1 μM TSA delay *D. discoideum* development, time to mound stage is shown in graph on left and complete development shown in graph on right. Each column represents the mean ± the standard error of the mean of three biological replicates performed in triplicate. **: $p<0.001$, *: $p<0.05$, ns: not significant, as assessed by one-way ANOVA analysis with Dunnett's multiple comparisons test. **d**, Treatment with 1 μM TSA and 10 μM BIX 01294 had no effect on *D. discoideum* fruiting body diameter while treatment with 5 μM sinefungin caused a reduction in fruiting body diameter. Fruiting body diameter correlates with the number of cells in each multicellular organism (*8, 9*). This graph represents the mean ± the standard error of the mean of three replicates performed with quantification of 5-10 fruiting bodies per replicate. Ns: not significant, *:p<0.05 as assessed by one-way ANOVA analysis with Dunnett's multiple comparisons test.

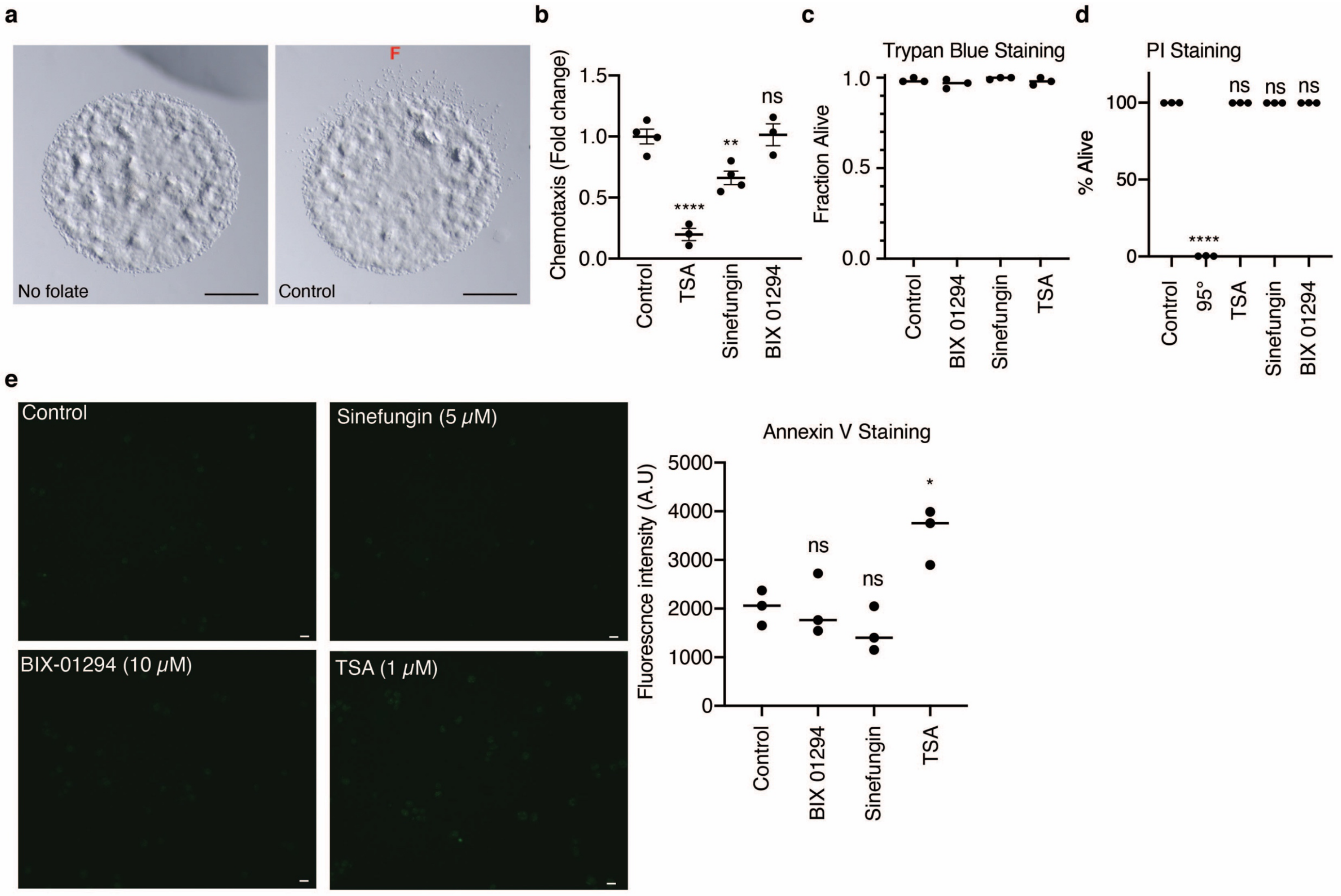


**Extended Data Fig. 8. Chemical inhibitors of chromatin modifications effect on chemotaxis and apoptosis**

**a,b**, Treatment with 10 μM BIX 01294 has no effect on *D. discoideum* chemotaxis ability while treatment with 5 μM sinefungin and 1 μM TSA inhibit chemotaxis. Representative images of control cells with or without 250 μM folate (red F on representative pictures) are displayed. The bar graph to the right represents the mean ± the standard error of the mean of three or four biological replicates performed in triplicate. The number of cells which migrated out of the spot where the *D. discoideum* were initially placed was counted in the 30˚ segment towards the 250 μM folate. Ns: not significant, ****:p<0.0001, **:p<0.001 as assessed by one-way ANOVA analysis. Scale bars are 100 microns. **c**, Treatment with 10 μM BIX 01294, 5 μM sinefungin, and 1 μM TSA have no effect on cell death as assessed by trypan blue staining. **d**, Treatment with 10 μM BIX 01294, 5 μM sinefungin, and 1 μM TSA have no effect on cell death as assessed by PI staining. 95°C heat shock for 50 seconds is sufficient to induce cell death. Each column represents the mean ± the standard deviation of three biological replicates. Ns: not significant, ****:p<0.0001 as assessed by one-way ANOVA analysis. **e**, Treatment with 10 μM BIX 01294 and 5 μM sinefungin have no effect on cell death as assessed by annexin V staining while treatment with 1 μM TSA increases annexin V staining. Representative images are displayed on the left while the graph on the right represents the mean ± the standard error of the mean of three biological replicates. Ns: not significant, *:$p<0.05$ as assessed by t-test.

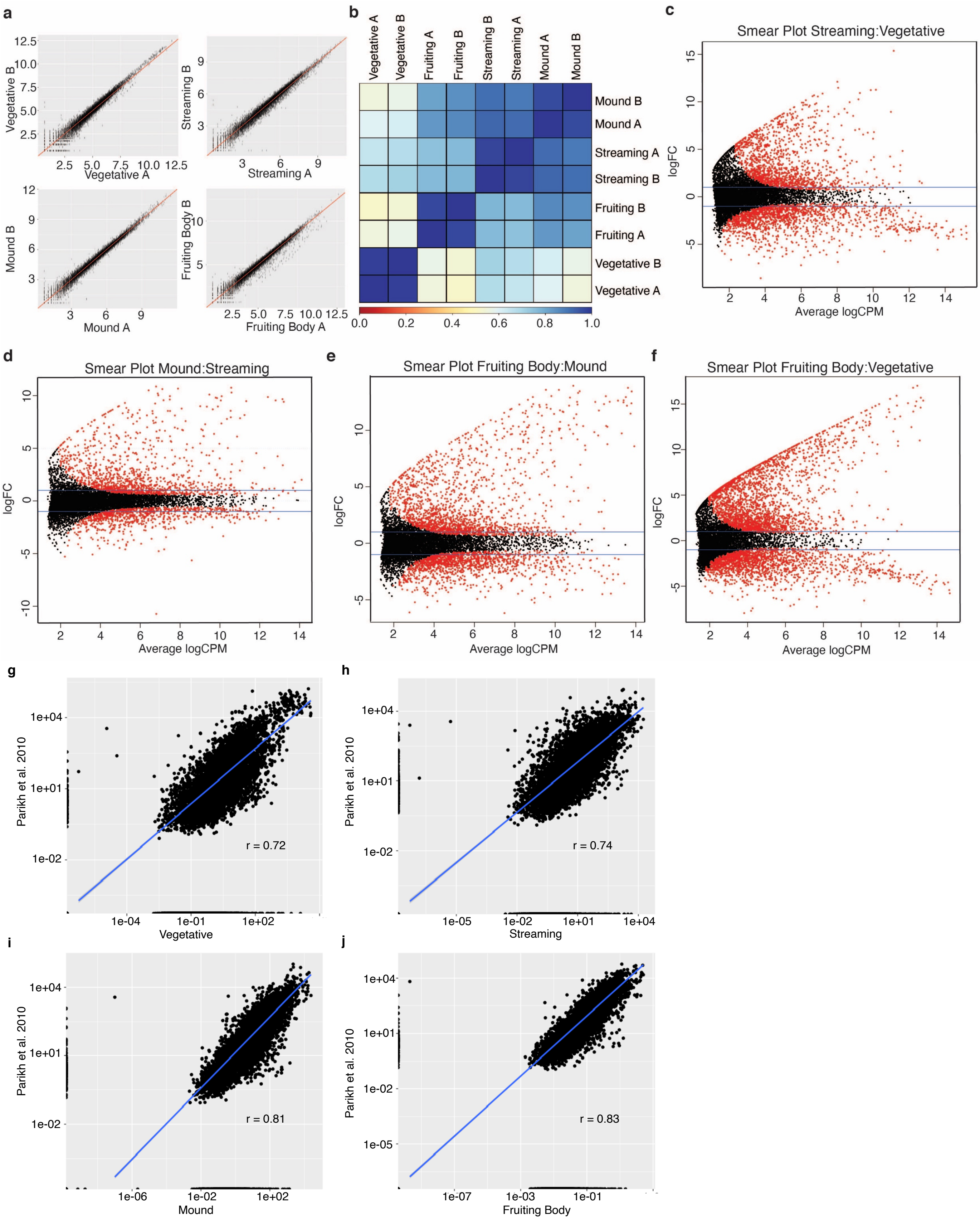
a
b
c
Smear Plot Streaming:Vegetative
d
Smear Plot Mound:Streaming
e
Smear Plot Fruiting Body:Mound
f
Smear Plot Fruiting Body:Vegetative
Average logCPM
logFC
g
h
i
j
Parikh et al. 2010
Vegetative
Streaming
Mound
Fruiting Body
r = 0.72
r = 0.74
r = 0.81
r = 0.83

**Extended Data Fig. 9 RNAseq analysis display a high degree of reproducibility between replicates and can distinguish developmental stages from each other**
**a**, A linear correlation between biological replicates is shown for each developmental stage. $r^2 > 0.95$ for each replicate. **b**, A heat map correlation of different RNAseq samples demonstrates the high degree of reproducibility and the differences between the different stages as well as unicellular compared to multicellular stages. **c-f,** A Bland-Altman plot of RNAseq datasets reveals genes which are upregulated and downregulated between individual stages of *D. discoideum*. Red dots represent significant differentially expressed genes, black dots represent not significant expressed genes. **g-j**, Comparison of bulk RNAseq expression to previously published RNAseq data (*27*) reveals a high degree of correlation.

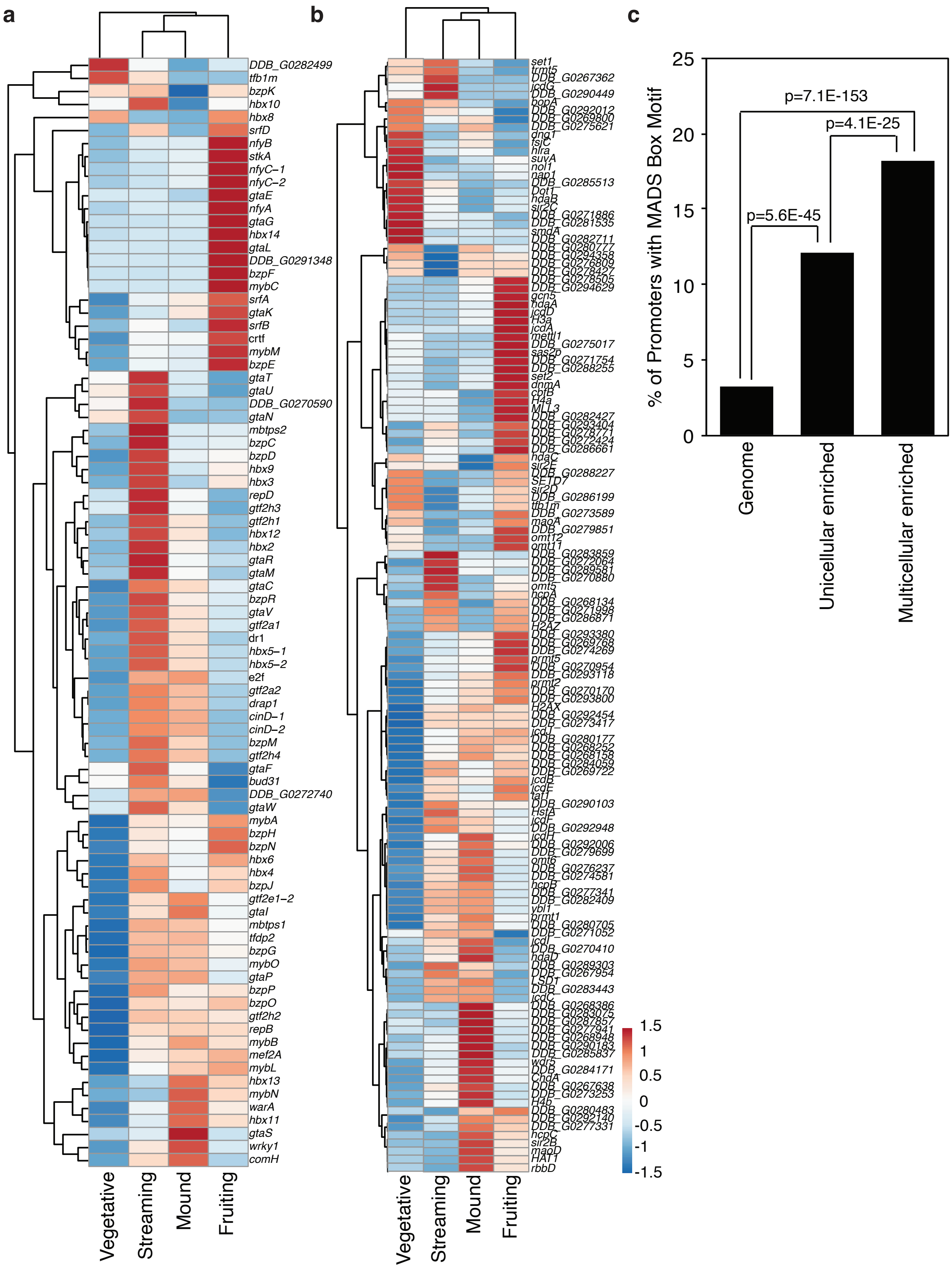


**Extended Data Fig. 10 Heat maps of a, transcription factors and b, chromatin modifying enzymes reveal stage specific expression patterns of several global regulators by RNAseq analysis. c, MADS box binding motif is significantly enriched in multicellular enriched**

**expressed genes relative to its occurrence in the genome and unicellular enriched expressed genes. P values were calculated by hypergeometric probability.**

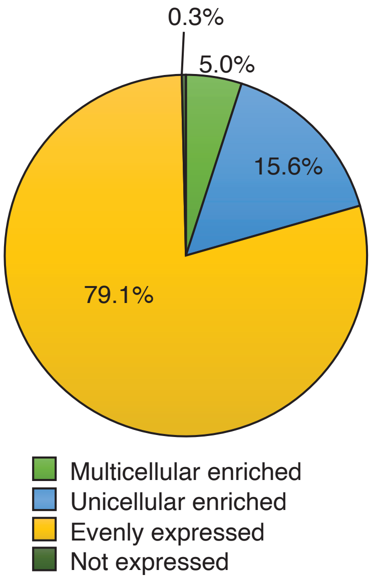


**Extended Data Fig. 11 Syntenic analysis of genes in *D. discoideum, S. pombe,* and *C. elegans* reveals unicellular or multicellular enrichment**

A pie-chart breaks down whether the 2473 conserved genes are expressed in both unicellular and multicellular stages (yellow; 79.1%), enriched expression in the unicellular stage (blue; 15.6%), enriched expression in the multicellular stages (light green; 5%) or not expressed (dark green; 0.3%).

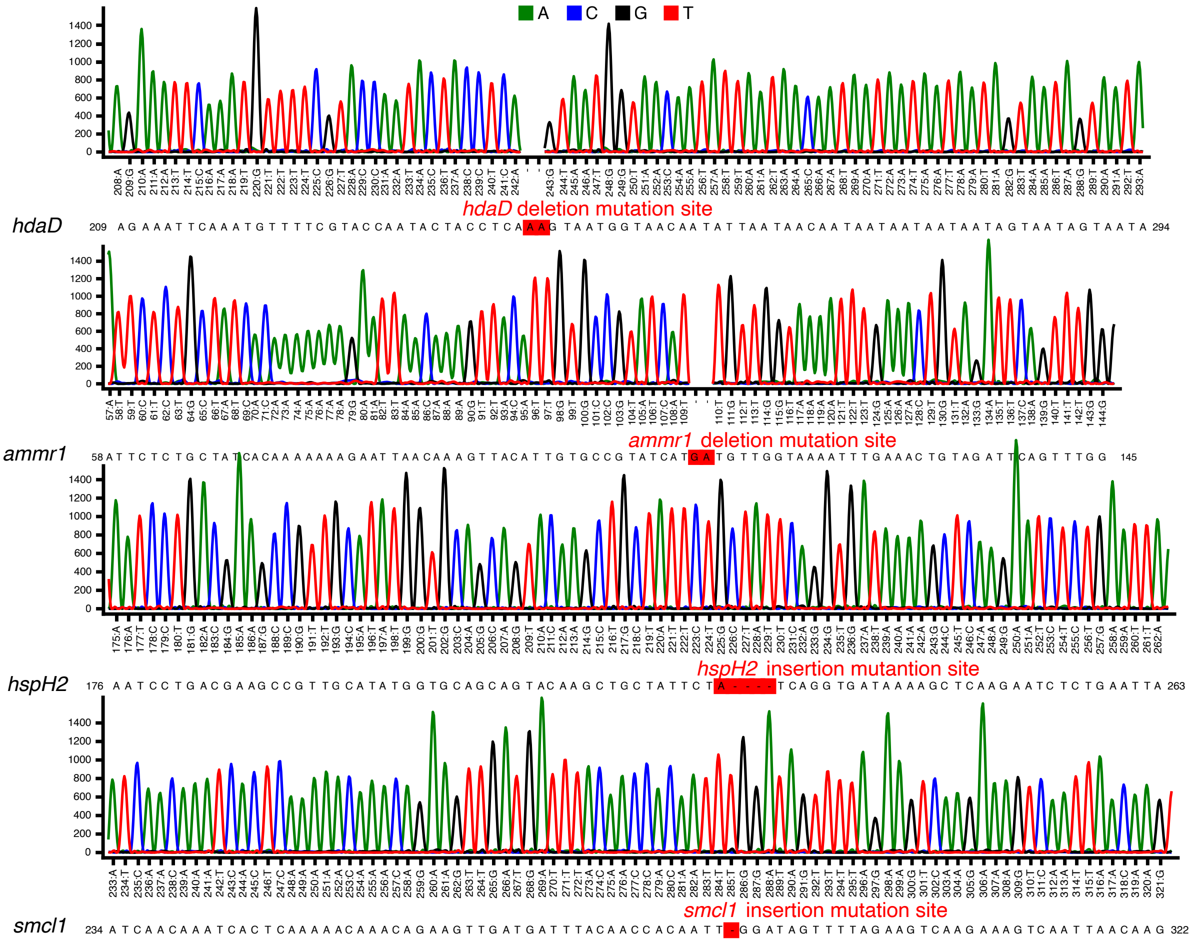


**Extended Data Fig. 12 Sequencing of deletion strains**
Each mutant strain was validated by sanger sequencing.

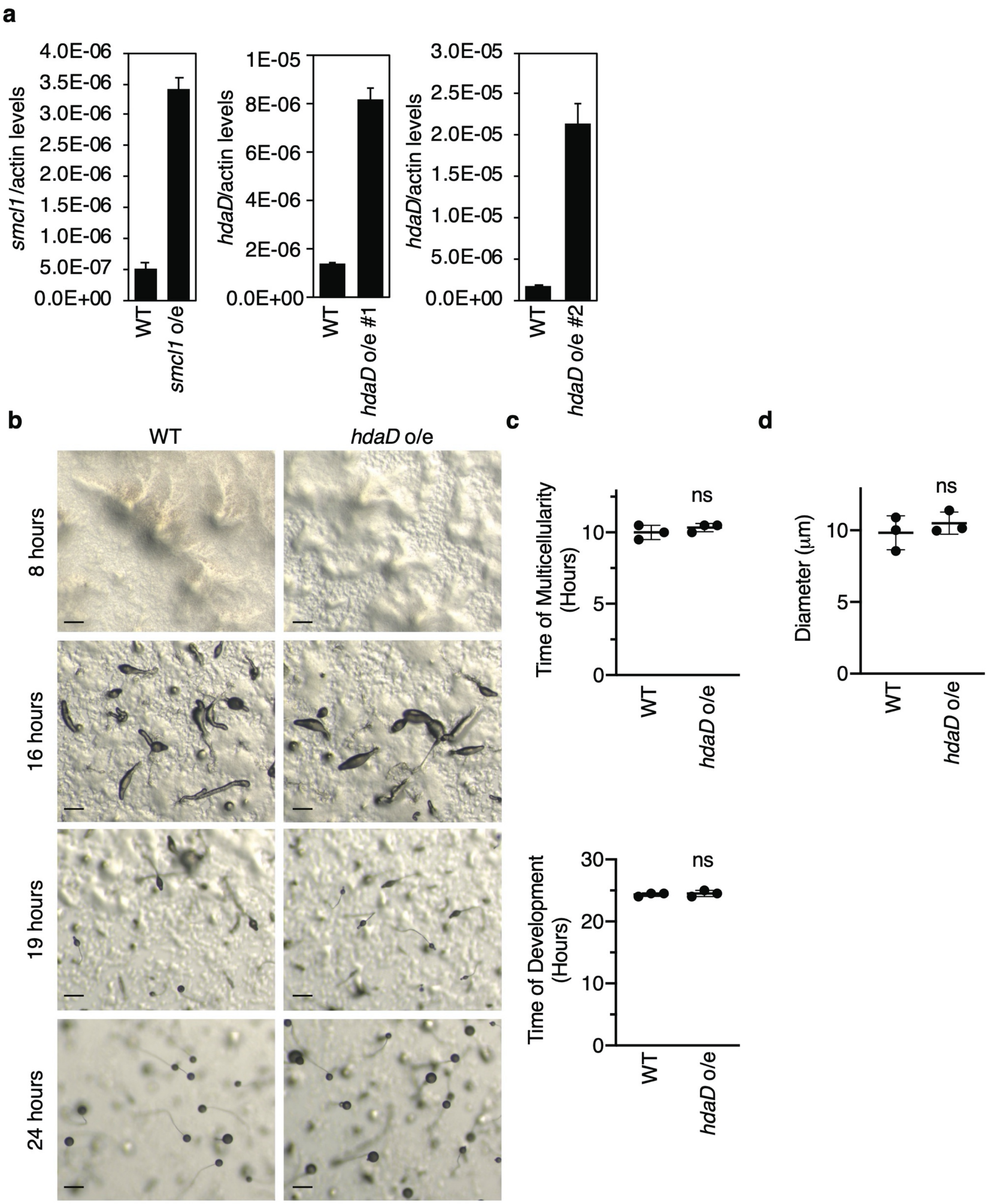


**Extended Data Fig. 13 *hdaD* overexpression has no effect on multicellularity**

**a**, Real time RT PCR confirms overexpression of *smcl1* and *hdaD*. Each bar represents the mean ± the standard deviation of 3 replicates. **b**, Representative images of *hdaD* overexpression reveals similar developmental timing to WT *D. discoideum*. Scale bars are 20 microns. **c**, Overexpression of *hdaD* causes no effect on multicellularity (upper graph) and total development (lower graph). This graph represents the mean ± the standard error of the mean of three independent biological replicates performed in triplicate. Ns: not significant, as assessed by unpaired t test. **d**, Overexpression of *hdaD* has no effect on the diameter of *D. discoideum* fruiting bodies. This graph represents mean ± the standard error of the mean of three independent biological replicates performed once. Ns: not significant, as assessed by unpaired t test.

**Table S1 ATACseq annotated enriched peak regions in unicellular and multicellular stages of *D. discoideum***

**Table S2 ChIPseq differential binding sites between unicellular versus multicellular stages of *D. discoideum***

**Table S3 ChIPseq annotated peak regions for H3K4me1, H3K4me3, H3K27ac, and H3K36me3**

**Table S4 RNAseq differentially expressed genes and gene ontology analysis**

**Table S5 Correlation of accessibility and gene expression**

**Table S6 Ortholog pairs among *D. discoideum, C. elegans* and *S. pombe***

**Table S7 Syntenic gene list**

**Table S8 Quality control summary for each sequencing experiment**
